# KMT2E recruitment by H3 serotonylation regulates neurodevelopmental chromatin dynamics

**DOI:** 10.64898/2026.09.20.752984

**Authors:** Jennifer C O’Chan, Benjamin H. Weekley, Celi Yang, Min Chen, Ashley M. Cunningham, Sohini Dutta, Winnie Chen, Rasika R. Iyer, Newaz I. Ahmed, Bulent Cetin, Cari A. Sagum, Bradley J. Lukasak, Erdene Baljinnyam, Zoé Christenson Wick, Vishwendra Patel, Emma Andraka, Aarthi Ramakrishnan, Christopher Peralta, Li Shen, Henrik Molina, Tristan Shuman, Robert D. Blitzer, Tom W. Muir, Mark T. Bedford, Samuele G. Marro, Haitao Li, Ian Maze

**Author notes:** **Corresponding Author:** Ian Maze. Denotes Equal contribution (Jennifer C O’Chan, Benjamin H. Weekley, Celi Yang).

## Abstract

Histone H3 Gln 5 serotonylation (H3Q5ser) is a recently identified epigenetic modification in brain that modulates *reader* interactions with adjacent H3 Lys 4 trimethylation (H3K4me3) to promote transcriptional permissiveness^1,2^. However, whether H3K4me3Q5ser and its associated binding proteins regulate neurodevelopmental gene expression programs remains unknown. Here, we identified the catalytically inactive Lysine methyltransferase 2E (KMT2E) as a *reader* of combinatorial H3K4me3Q5ser. KMT2E preferentially binds H3K4me3Q5ser over H3K4me3 alone, and enriches at broad chromatin domains marking actively transcribed neurodevelopmental loci. Notably, heterozygous variants in *KMT2E* have been implicated in ODLURO syndrome, a recently characterized neurodevelopmental disorder (NDD)^3,4^. To identify the molecular mechanisms underlying ODLURO syndrome, we generated a *Kmt2e* transgenic mouse model that reproduces behavioral, physiological, and cellular endophenotypes associated with this and other NDDs. Furthermore, we observed that KMT2E mediates these effects by recruiting the NCoR/HDAC3 repressor complex to H3K4me3Q5ser-marked loci to restrict spreading of co-localized H3 Lys 9 acetylation (H3K9ac). Inhibition of aberrant H3K9ac spreading was sufficient to rescue transcriptional dysregulation in *Kmt2e* haploinsufficient neurons. These findings thus establish KMT2E as a critical reader of H3 serotonylation during neurodevelopment and provide mechanistic insights into the pathogenesis of ODLURO syndrome.

## INTRODUCTION

Serotonin (5-hydroxytryptamine, 5-HT) is a biogenic amine that is critical for nervous system development via its regulation of neuronal proliferation, migration, and circuit formation through classic receptor-mediated signaling^5,6^. Given its regulatory roles in guiding synaptogenic processes, perturbations to 5-HTergic signaling during early developmental periods can produce long-lasting effects on brain organization and function, as evidenced in multiple neurodevelopmental disorders (NDDs)^7,8^. Thus, understanding the full complement of mechanisms through which 5-HT shapes the developing brain is crucial to understanding the etiologies of NDDs.

Recently, a novel receptor-independent role for 5-HT in the nucleus, termed histone serotonylation, has been described by our group and others. In the nucleus, 5-HT can be covalently attached to histone H3 at Gln 5 (H3Q5ser) by the enzyme transglutaminase 2 (TG2) within chromatin-bound nucleosomes. H3Q5ser often co-occurs with adjacent Lys 4 trimethylation (H3K4me3), thus establishing the combinatorial H3K4me3Q5ser mark^1^. In the context of H3K4me3, H3Q5ser modulates the binding of H3K4me3 *reader* proteins to influence downstream transcription^1,2,9–16^. Given that dysregulation of H3K4me3 *writers*, *erasers*, and *readers* have been shown to underlie several NDDs^17,18^, understanding how this post-translational modification (PTM) is regulated by H3Q5ser during neurodevelopment represents an important, yet previously uncharacterized, intersection between canonical 5-HTergic signaling and epigenetic dysregulation. Previously, a subset of H3K4me3 peaks spanning continuous ‘broad’ genomic domains (>4 kb, up to 60 kb) has been described^19,20^. These broad domains are enriched at cell-type-specific loci, and have been proposed to regulate transcriptional permissiveness of cognate genes, including those important for synaptic signaling during brain development. Furthermore, dysregulation of these broad H3K4me3 domains has been observed in postmortem prefrontal cortical tissues from individuals with autism spectrum disorder, where H3K4me3 has been found to aberrantly spread into gene bodies^21,22^. However, the mechanisms regulating such spreading, and the impact of these phenomena on neurodevelopmental transcriptional trajectories, remain unknown.

Here, we identified Lysine methyltransferase 2E (*KMT2E*, also termed mixed lineage leukemia 5 (MLL5)) as a *reader* of H3K4me3Q5ser in the developing brain. Unlike other members of the KMT2 family, KMT2E lacks intrinsic methyltransferase activity. Instead, it has been shown to function primarily through its plant homeodomain (PHD) to bind H3K4me3, an interaction that we found was enhanced by the presence of H3Q5ser^23–25^. In *S. cerevisiae* and *D. melanogaster*, the *KMT2E* orthologs *SET3/SET4* and *UpSET*, respectively, are known to recruit histone deacetylase (HDAC) complexes to active gene loci^26–29^. However, the functional consequences of this evolutionarily conserved mechanism has yet to be characterized in mammals, especially in brain. Furthermore, loss of *UpSET* in *Drosophila* results in aberrant ‘spreading’ of both H3K9ac and H3K4me3 past their normal boundaries at active TSSs^26^. In humans, O’Donnell-Luria-Rodan (ODLURO) syndrome, an autosomal dominant NDD that is characterized by symptoms including developmental delay, intellectual disability, hypotonia, and epilepsy, arises from heterozygous variants within the *KMT2E* locus^3,4,30^. These variants include missense and protein-truncating mutations, copy number variants, and chromosomal microdeletions that encompass the *KMT2E* locus. Despite ODLURO syndrome stemming from a monogenic cause, the mechanisms underlying its pathogenesis remain unknown. Using a translationally relevant *Kmt2e* transgenic mouse model, we demonstrate that KMT2E binds H3K4me3Q5ser at gene loci important for neurodevelopmental processes, where it subsequently recruits the NCoR/HDAC3 complex to mediate histone PTM spreading and downstream transcriptional regulation. Together, these findings comprehensively establish a mechanistic basis for ODLURO syndrome, and position histone serotonylation as a critical node in the regulation of neurodevelopmental gene programs.

## RESULTS

### KMT2E is a reader of H3 serotonylation

To assess potential roles for H3 serotonylation in developing brain, we performed ChIP-sequencing using a validated H3K4me3Q5ser antibody^1^ on embryonic day (E) 12.5 mouse brain, a timepoint that follows dorsal raphe nucleus specification^6^ and corresponds with increased H3K4me3Q5ser levels. We identified 20,267 peaks genome-wide, predominantly near transcriptional start sites (TSSs), consistent with previous reports^1,10^ (**Extended Data Fig. 1a)**. These findings suggested that H3 serotonylation may play widespread and functionally significant roles in regulating transcription during brain development. As prior work demonstrated distinct transcriptional features associated with H3K4me3 peak breadth^19,20^, we next explored whether H3 serotonylation contributes to these domains by classifying H3K4me3Q5ser peaks as narrow (<5 kb) *vs.* broad (≥5 kb). Indeed, H3K4me3Q5ser was found to associate with both narrow and broad H3K4me3Q5ser domains, and co-localized with corresponding narrow *vs.* broad H3K4me3 and H3K9ac signals in E12.5 brain tissues^31^ (**Fig. 1a**). Such patterns were consistent with potential coordinated regulation of peak breadth across the three PTMs, which have long been associated with TSSs of active genes^32^. Consistent with published work demonstrating that broad H3K4me3 domains mark highly expressed, cell-type specific genes^20^, broad H3K4me3Q5ser loci similarly exhibited significantly higher expression *vs.* narrow peak loci (**Fig. 1b**) and were enriched for ontology terms related to neurodevelopmental processes including nervous system development, axon guidance, and neuron differentiation (**Fig. 1c**). In contrast, narrow peak-associated genes were found to be enriched for housekeeping processes including rRNA processing, cellular respiration, and aerobic electron transport (**Fig. 1d**). In total, these findings supported the notion that broad H3K4me3Q5ser domains specifically mark actively transcribed neurodevelopmental gene expression programs during embryonic brain development.

**Figure 1.**
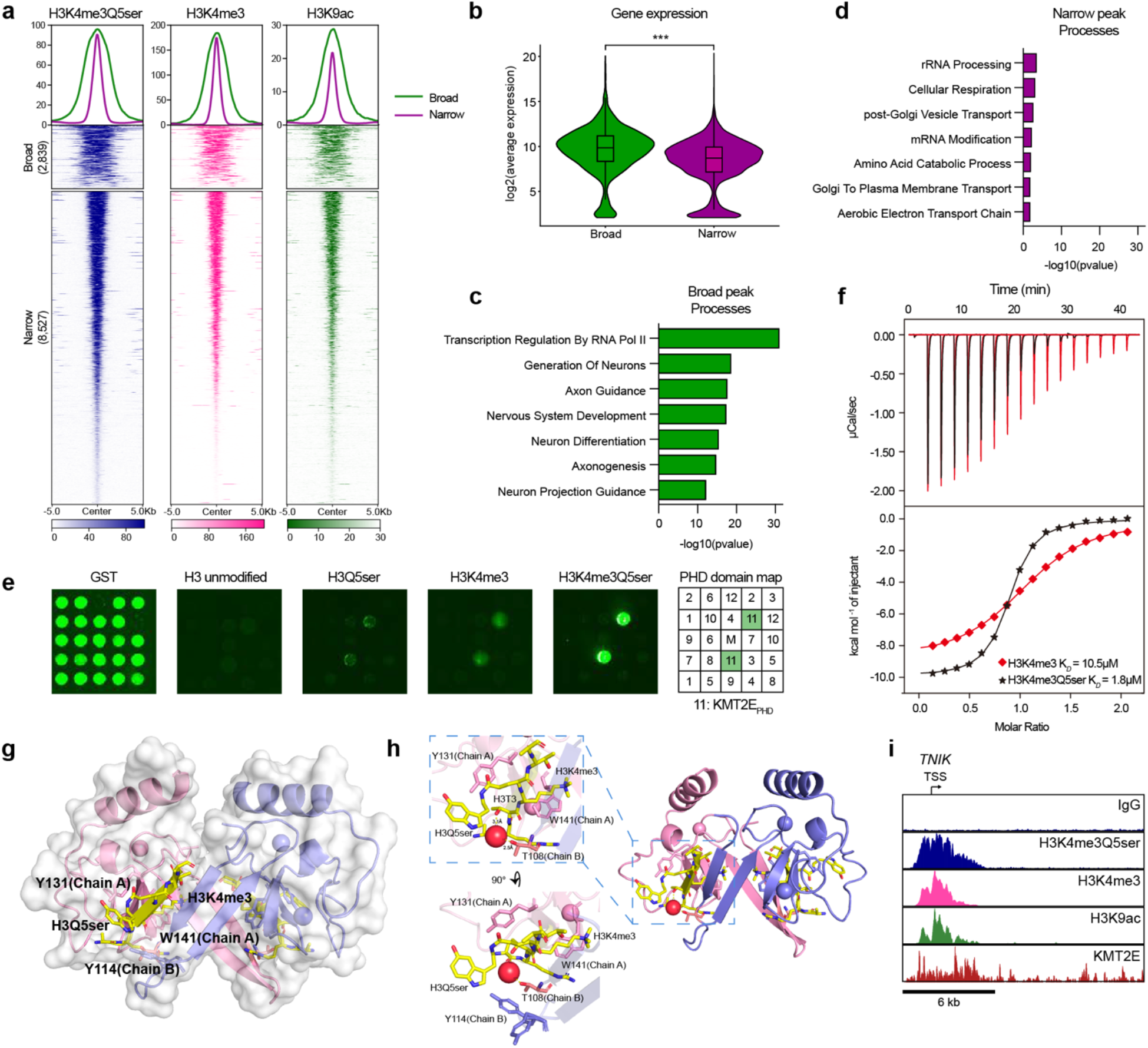
KMT2E is a *reader* of H3K4me3Q5ser. (**a**) ChIP-seq heatmaps and profiles for broad (≥ 5k bp) vs. narrow (<5k bp) H3K4me3Q5ser, H3K4me3, and H3K9ac peaks at TSSs for genes expressed in E12.5 mouse brain. (**b**) Gene expression levels associated with broad *vs.* narrow H3K4me3Q5ser peaks in E12.5 brain (*n* = 4). (**c-d**) Significantly enriched pathways from GO analysis of (**c**) broad and (**d**) narrow H3K4me3Q5ser peaks (adjusted *p-value* < 0.05). (**e**) Protein *reader* domain microarray of select PHDs following binding of biotinylated H3_1-10_ peptides (unmodified, H3Q5ser, H3K4me3, H3K4me3Q5ser), demonstrating selectively enhanced interactions with KMT2E_PHD_ (run in duplicate); anti-GST loading controls are provided (left). Labels for the full panel can be found in **Extended Data Figure 1**. (**f**) Titration and fitting curves of peptides titrated into KMT2E_PHD_. K_D_ values are provided. See **Extended Data Table 1** for ITC statistics. (**g**) Structure of the KMT2E_PHD_-H3K4me3Q5ser complex, with enlarged inset shown in **(h)**. Ribbon shows detailed construction of KMT2E_PHD_, with key amino acid residues highlighted in Chain A (pink) *vs.* Chain B (purple). Yellow sticks show H3 residues. See **Extended Data Table 2** for x-ray crystallography data collection and refinement statistics. (**i**) Representative genome browser tracks of broad H3K4me3Q5ser, H3K4me3, and H3K9ac peaks at a KMT2E enriched locus (*n* = 4) in mouse brain (*vs.* IgG). For (a) and (i), H3K4me3Q5ser (*n* = 4); data for H3K9ac and H3K4me3 from Gorkin et al. (2020)^31^; *n* = 2 per PTM); for (i), KMT2E (*n* = 3).

Next, to identify potential mechanisms underlying broad H3K4me3Q5ser domain regulation, we employed a high-throughput histone *reader* domain array in which biotinylated histone H3 tail peptides (unmodified, H3Q5ser, H3K4me3, or H3K4me3Q5ser) were screened against 250 purified *reader* domains^33^ (**Extended Data Fig. 1b-f**). Using this approach, we identified several domains which bound H3K4me3, H3Q5ser, and H3K4me3Q5ser. Of the 19 domains which were identified to bind H3K4me3Q5ser, the majority showed enhanced binding in comparison to H3K4me3 alone. Notably, 8 of these 19 domains exist within chromatin regulatory proteins that have previously been implicated in NDDs and related disorders^34–40^. Two domains that displayed the greatest level of enhanced binding to H3K4me3Q5ser relative to H3K4me3 or H3Q5ser alone were the chromo domain of CHD1 and the PHD finger of KMT2E (KMT2E_PHD_) (**Fig. 1e, Extended Data Fig. 1b-f**). Previously, we reported that the dissociation constant (K_d_) for CHD1’s chromo domain is enhanced ∼2.25 fold in the presence of H3K4me3Q5ser relative to H3K4me3 alone (∼ 12 μM *vs.* 5 μM)^10^. For the KMT2E_PHD_ domain, we observed a K_d_ of 10.5 μM for H3K4me3 (consistent with prior reports^23^) *vs.* 1.8 μM for H3K4me3Q5ser, representing a 5.8-fold enhancement in binding (**Fig. 1f**). Due to such greater enhancement and more robust binding of KMT2E to H3K4me3Q5ser *vs.* CHD1, as well as previously documented evidence that KMT2E’s orthologues play important roles in broad domain regulation^26^, we chose to focus on KMT2E for the remainder of this study. Notably, this enhanced binding was not observed for the PHD fingers of other KMT2/MLL family members (**Fig. 1e, Extended Data Fig. 1g**). To validate this interaction, we next expressed and purified recombinant His-tagged KMT2E_PHD-SET_ domains and confirmed preferential binding to H3K4me3Q5ser via *in vitro* peptide pulldown assays (**Extended Data Fig. 2a)**. Such enhanced binding was independently corroborated by peptide pulldown assays from HeLa cell nuclear extracts followed by western blotting for endogenous KMT2E (**Extended Data Fig. 2b)**.

In order to define the molecular basis of this enhanced interaction, we further solved the X-ray crystal structure of KMT2E_PHD_ in complex with an H3K4me3Q5ser peptide, where full electron density was traced across the binding interface (**Extended Data Fig. 2c**). Structural analysis revealed that the Trp (W) 141 residue within Chain A engages H3K4me3 through a cation-π bond interaction, while Tyr (Y) 131 of Chain A forms a parallel-displaced amide-π interaction with the side-chain amide of H3Q5ser, with the aromatic ring positioned nearly coplanar (∼8°) to the amide plane at a distance of 4.3–4.6 Å, collectively stabilizing recognition of the dually modified H3 tail. (**Fig. 1g-h**). Importantly, mutating KMT2E_PHD_ Y131 to Ala (A) reduced its affinity to H3K4me3Q5ser by ∼2-fold (**Extended Data Fig. 2d**). In addition, we observed that recognition of H3Q5ser occurs via hydrogen bonding, involving a direct hydrogen bond between the 5-hydroxyl group of the serotonin moiety and the backbone carbonyl oxygen of Arg (R) 151 in Chain B, with an O–O distance of 3.2 Å, contributing to stabilization of the serotonylated glutamine within the binding pocket. (**Fig. 1g-h**). There is apparent stabilization of the peptide binding via a water-mediated hydrogen bonding network, in which a water molecule bridges H3 threonine (Thr/T) 3 to T108 of Chain B at 3.1/2.5 Å respectively (**Fig. 1h**).Together, these data indicated that H3Q5ser likely enhances recruitment of KMT2E to H3K4me3 to regulate gene expression via modulation of histone PTM breadth. We confirmed this relationship using KMT2E CUT&RUN-sequencing from mouse brain, which demonstrated that KMT2E occupancy occurs at broad H3K4me3Q5ser, H3K4me3, and H3K9ac domains (e.g., at the *TNIK* locus; **Fig. 1i**).

### Loss of Kmt2e recapitulates features of ODLURO syndrome in mice

Heterozygous variants in *KMT2E*, the majority of which are predicted to result in loss of functional KMT2E protein, have been described in patients with ODLURO syndrome^3,4,30^. Thus, to investigate potential deleterious consequences of *KMT2E* haploinsufficiency during neurodevelopment and to establish a mouse model of ODLURO syndrome with construct, face, and predictive validity, we generated *Kmt2e* constitutive knockout mice by introducing a frameshift mutation in coding exon 2, which produces a premature stop codon (**Fig. 2a, Extended Data Fig. 3**). Loss of *Kmt2e* gene expression and genome-wide chromatin occupancy in brain tissues were confirmed in *Kmt2e^-/-^* (KO) animals by qPCR and CUT&RUN-sequencing, respectively (**Fig. 2b-c**). Consistent with prior reports of embryonic lethality in an independent line of *Kmt2e^-/-^*mice^41^, we observed significant sub-viability of homozygous KO animals at weaning relative to expected Mendelian ratios (**Fig. 2d**), thus underscoring essential roles for *Kmt2e* during development.

**Figure 2.**
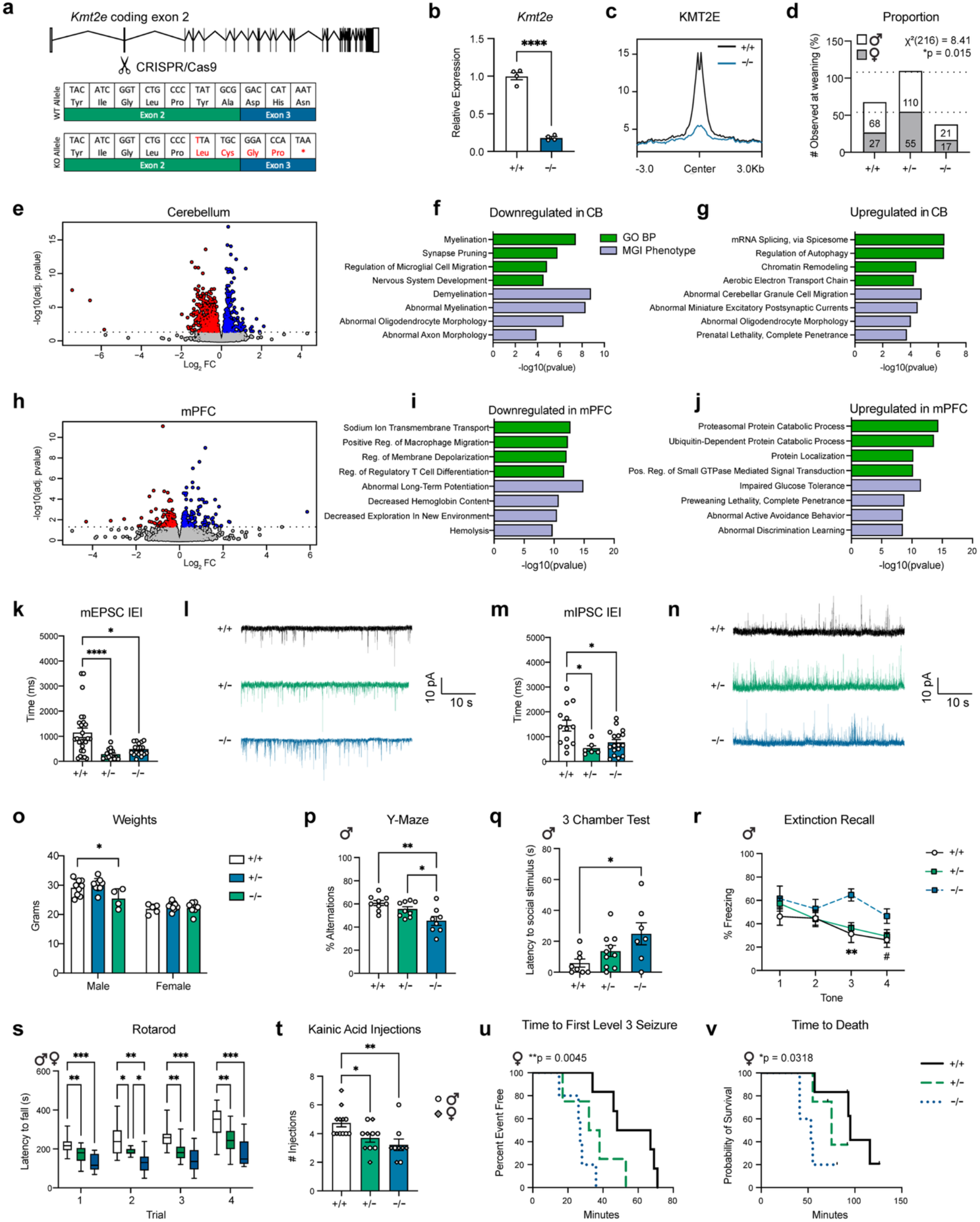
KMT2E loss-of-function disrupts transcriptional, physiological, and behavioral outcomes in mice. (**a**) CRISPR/Cas9-mediated targeting of coding exon 2 of the *Kmt2e* locus to generate a mutant KO allele (top), resulting in a 1 bp frameshift insertion that introduces a premature stop codon (bottom). (**b**) *Kmt2e* expression is reduced in KO brain tissues. Two-sided unpaired Student’s t-test: \*\*\*\**p* < 0.0001; *n* = 4/genotype. (**c**) CUT&RUN-seq demonstrating that KMT2E genomic occupancy is reduced in KO brain. *n* = 3/genotype. (**d**) Fewer Kmt2e^-/-^ mice are observed at postnatal day 21 compared to expected Mendelian ratios (χ² (216) = 8.41; *p* = 0.015). The number of males and females per genotype are presented. (**e, h**) Volcano plots displaying DEGs (*padj* < 0.05) between WT *vs.* KO mice at P14 for (**e**) cerebellum (CB) and (**h**) mPFC. *n* = 4/genotype/brain region. Pathway analysis for differentially expressed genes that are (**f**) downregulated in CB, (**g**) upregulated in CB, (**i**) downregulated in mPFC, and (**j**) upregulated in mPFC. (**k**) Cell means and (**l**) representative traces for interevent intervals (IEI) of mEPSCs for WT (*n* = 26), HET (*n* = 19), and KO (*n* = 24) cerebellar granule neurons. Kruskal-Wallis test: *p* = 0.0001; Dunn’s *post-hoc*: WT *vs.* HET, \*\*\*\**p* < 0.0001; WT *vs.* KO, \**p* = 0.0412. (**m**) Cell means and (**n**) representative traces for IEI of mIPSCs for WT (*n* = 13), HET (*n* = 7), and KO (*n* = 17) cerebellar granule neurons. Kruskal-Wallis test: *p* = 0.008; Dunn’s *post-hoc*: WT *vs.* HET, \**p* = 0.0119; WT *vs.* KO, \**p* = 0.0316. (**o**) Weights for adult male (WT, *n* = 9; HET, *n* = 9; KO, *n* = 4) and female (WT, *n* = 9; HET, *n* = 5; KO, *n* = 7) mice. Two-way ANOVA: effect of genotype, *p* = 0.0351; sex, *p* < 0.0001; interaction, *p* = 0.0395. Tukey’s *post-hoc*: male WT *vs.* KO, \**p* = 0.0158. (**p**) KO (*n* = 8) mice displayed reduced spontaneous alternations compared to WT (*n* = 9) and HET (*n* = 9) mice on the Y-maze. One-way ANOVA: *p* = 0.0026. Tukey’s *post-hoc*: WT *vs*. KO, *\*p* = 0.045; HET *vs.* KO, \*\**p* = 0.002. (**q**) KO (*n* = 7) mice exhibited increased latency to approach a novel social stimulus compared to WT (*n =* 8) and HET (*n* = 10) mice. One-way ANOVA: *p* = 0.032. Tukey’s *post-hoc*: WT *vs.* KO, *\*p* = 0.0251. (**r**) KO (*n* = 8) males displayed reduced extinction recall following auditory fear conditioning compared to WT (*n* = 11) and HET (*n* = 21) mice. Two-way RM ANOVA: time: *p* < 0.0001; genotype, *p* = 0.0616. Dunn’s *post-hoc*: tone_3_: WT *vs.* KO, \*\**p* = 0.0033; tone_4_: WT *vs.* KO, #*p* = 0.0631. (**s**) HET (*n* = 13) and KO (*n* = 9) mice displayed impaired motor coordination and learning on the rotarod test compared to WT (*n* = 15) mice. Two-way RM ANOVA: trial: *p* < 0.0001; genotype, *p* < 0.0001. Tukey’s *post-hoc*: \*\*\**p* < 0.001, \*\**p* < 0.01, \**p* < 0.05. Box plot shows median (center line), interquartile range (box), and minimum-to-maximum values (whiskers). (**t**) HET (*n* = 10) and KO (*n* = 9) mice required fewer kainic acid injections to produce a level 3 seizure compared to WT (*n* = 12). One-way ANOVA: *p* = 0.0063. Dunn’s *post-hoc*: WT *vs.* HET, *\*p* = 0.0462; WT *vs.* KO, *\*\*p* = 0.0045. (**u-v**) Survival curves showing a significant effect of genotype using the log-rank test for time to (**u**) first level 3 seizure (χ² = 10.83, df = 2, *p* = 0.0012) and (**v**) death (χ² = 6.894, df = 2, *p* = 0.0318) in female mice. For u-v: WT (*n* = 6), HET (*n* = 4), KO (*n* = 5). Bar plots are presented as mean ± SEM. See **Source Data** file for number used to generate bar plots.

To confirm that *Kmt2e* loss-of-function alters actively transcribed neurodevelopmental gene programs, we then performed bulk mRNA-sequencing from mouse cerebellum and medial prefrontal cortex (mPFC) at postnatal day (P) 14, a timepoint associated with active synaptogenesis and circuit maturation in both regions^42,43^. These regions were selected based upon their established involvement in autism- and NDD-related pathologies^44,45^. In ODLURO patients, cerebellar dysplasia and cerebral atrophy have been documented, further supporting selection of these brain regions^3,4^. While both cerebellum and mPFC exhibited significant transcriptional alterations in *Kmt2e^-/-^* mice, the cerebellum displayed a greater number of differentially expressed genes (DEGs), with downregulated genes being enriched for processes associated with myelination, synaptic pruning, and nervous system development, and upregulated genes displaying enrichment for ontologies related to chromatin remodeling, mRNA splicing, and abnormal miniature excitatory postsynaptic currents (mEPSCs) (**Fig. 2e-g**). In mPFC, downregulated genes were enriched for synaptic transmission-related processes including sodium ion transport, membrane depolarization, and long-term potentiation, while upregulated genes were enriched for proteasomal and ubiquitin-dependent protein catabolic processes (**Fig. 2h-j**). Together, these data indicated that *Kmt2e* loss-of-function disrupts distinct neurodevelopmental gene programs in a brain region-specific manner.

To further determine how *Kmt2e* impacts these gene expression programs in brain, we examined transcription factors as potential co-regulators. ChEA analysis revealed distinct transcription factors underlying cerebellar gene expression, including CREB1 and CREM for upregulated DEGs, and OLIG2 and MBD3 for downregulated DEGs (**Extended Data Fig. 4a**). Given that ontology enrichments such as splicing, abnormal miniature postsynaptic currents, and myelination were observed for upregulated cerebellar DEGs, we then examined the functional consequences of *Kmt2e* loss on these measures. We identified significantly more differential splicing events in the cerebellum of *Kmt2e^-/-^* animals relative to mPFC, with skipped exon and retained intron events being most prominent (**Extended Data Fig. 4b**). Next, we examined the physiological consequences of *Kmt2e* KO in cerebellar granule neurons using patch-clamp electrophysiology. Both mEPSC and miniature inhibitory postsynaptic current (mIPSC) inter-event intervals were found to be significantly reduced in KO animals relative to WT controls, indicating heightened spontaneous synaptic activity with no significant differences observed in amplitude (**Fig. 2k-n, Extended Data Fig. 4c-d**). We additionally confirmed altered myelination in *Kmt2e^-/-^* animals by electron microscopy, with KO animals exhibiting a greater number of myelinated fibers with reduced myelin thickness, as indicated by increased g-ratios in comparison to WT mice (**Extended Data Fig. 4e-i**). In mPFC, assessments of cortical layer markers revealed disruption of deep cortical layers specifically, with altered expression of Layer V/VI markers (i.e., CTIP2 and TBR1) observed in *Kmt2e^-/-^* animals, while no differences were observed for the upper layer marker BRN1^46^ (**Extended Data Fig. 4j-o**). These findings are suggestive of layer-specific impairments of cortical lamination that recapitulate endophenotypes observed across numerous NDDs. Notably, disruption of CTIP2 and TBR1, markers of corticospinal motor and corticothalamic projection neurons, respectively, may contribute to motor and cognitive endophenotypes observed in ODLURO patients.

To next determine whether these molecular and cellular disruptions manifest as developmental and behavioral consequences, we assessed *Kmt2e^-/-^*animals across numerous paradigms relevant to ODLURO syndrome endophenotypes. Body weight assessments revealed differences by genotype, but not sex, at P14 (**Extended Data Fig. 5a**). Sex-dependent effects in weight emerged in adulthood, with male KO animals exhibiting significantly reduced body weights relative to WT controls (**Fig. 2o**). Given that ODLURO syndrome is more prevalent in males and displays sex-dependent symptom expression^3,4^ (e.g., features of autism predominating in males, and epilepsy in females), we continued to characterize sex-dependent phenotypic divergence in our model. We first examined autism-relevant behavioral endophenotypes, including working memory, social preference, and cognitive flexibility. On the Y-maze, we observed reduced spontaneous alternations in *Kmt2e* KO males, but not females (**Fig. 2p, Extended Data Fig. 5b**). In the three chamber social preference test, male KO animals exhibited increased latency to approach a novel social stimulus relative to a novel non-social stimulus, consistent with impaired social preferences previously reported in another Kmt2e loss-of-function model; no differences were observed in females (**Fig. 2q, Extended Data Fig. 5c-e**). In auditory fear conditioning, no significant differences in acquisition, recall, or extinction were observed; however, extinction recall was found to be impaired in KO males (**Fig. 2r, Extended Data Fig. 5f-l**). To assess motor phenotypes, rotarod testing revealed that *Kmt2e^+/-^* (HET) and KO animals of both sexes displayed impaired baseline motor performance on trial 1, which failed to improve across subsequent trials (**Fig. 2s**). Notably, open field testing revealed no differences in the total distance traveled in male KO mice (**Extended Data Fig. 5m**), suggesting that performance differences observed in non-motor behaviors are not likely due to altered locomotor activity, while rotarod deficits reflect specific impairments in coordination and motor learning. We also observed altered anxiety-like behavior selectively in HET animals of both sexes, evidenced by increased center time in the open field (additionally corroborated in the light-dark box task) (**Extended Data Fig. 5n-o**). Finally, as epilepsy is a key feature of ODLURO syndrome, we assessed seizure susceptibility using a kainic acid-induced seizure paradigm. HET and KO animals were found to require significantly fewer injections to reach a level 3 seizure (as defined by the Racine scale) relative to WT mice, indicating increased susceptibility (**Fig. 2t**). In particular, female KO animals exhibited reduced latency to first level 3 seizure and reduced survival relative to WT, with no significant effects observed in males (**Fig. 2u-v, Extended Data Fig. 5p-q**). Together, these data recapitulate sex variances in symptomatic distribution observed in ODLURO patients, supporting the utility of this model for investigating the molecular mechanisms underlying KMT2E function during neurodevelopment.

### KMT2E recruits the NCoR/HDAC3 complex to regulate H3K9ac spreading

Having established the transcriptional, physiological, and behavioral consequences of *Kmt2e* loss-of-function, we next sought to elucidate the molecular basis by which KMT2E mediates these effects. We first confirmed that KMT2E lacks intrinsic methyltransferase activity, as neither the SET domain alone, PHD finger, nor combined PHD+SET domains produced detectable H3K4 methyltransferase activity *in vitro* (**Extended Data Fig. 6a-b**). This is consistent with prior reports indicating evolutionary conservation of a catalytically dead SET domain across *KMT2E* orthologs due to mutations in the s-adenosylmethionine binding site (**Extended Data Fig. 6c**)^23,25^.

To then assess potential non-enzymatic mechanisms through which KMT2E recruitment to H3K4me3Q5ser regulates chromatin dynamics, we generated a CRISPR-Cas9 mediated knock-in HeLa-S3 cell line, where a C-terminal 3xFLAG tag on the endogenous full-length *KMT2E* was introduced. After confirming high editing efficiency and FLAG-tagged KMT2E expression (**Extended Data Fig. 6d-f**), we performed FLAG immunoprecipitations from *KMT2E*-3xFLAG *vs.* WT HeLa-S3 nuclear extracts, followed by liquid chromatography-mass spectrometry. Using this unbiased approach, we identified all four core subunits of the Nuclear Corepressor (NCoR)/HDAC3 complex (NCOR1, NCOR2, TBL1XR1, and HDAC3), alongside the H3K4me3 demethylase JARID1A (KDM5A) (**Fig. 3a**), as potential KMT2E-interacting proteins/complexes. These interactions were validated by KMT2E-FLAG co-immunoprecipitation followed by western blotting, with non-interacting proteins included to confirm specificity (**Fig. 3b**). NCOR1 co-immunoprecipitation revealed strong enrichment of KMT2E, suggesting a direct interaction within the NCoR complex, while HDAC3 co-immunoprecipitation showed only modest KMT2E enrichment, consistent with the known participation of HDAC3 in multiple chromatin modifying complexes^47^ (**Extended Data Fig. 6g**). As the NCoR/HDAC3 complex and JARID1A are established *erasers* of histone acetylation and H3K4me3^48^ respectively, these results suggested that KMT2E recruitment to H3K4me3Q5ser may function to restrict overall levels and spreading of these PTMs at active TSSs, as supported by the previous findings in *Drosophila*^26^.

**Figure 3.**
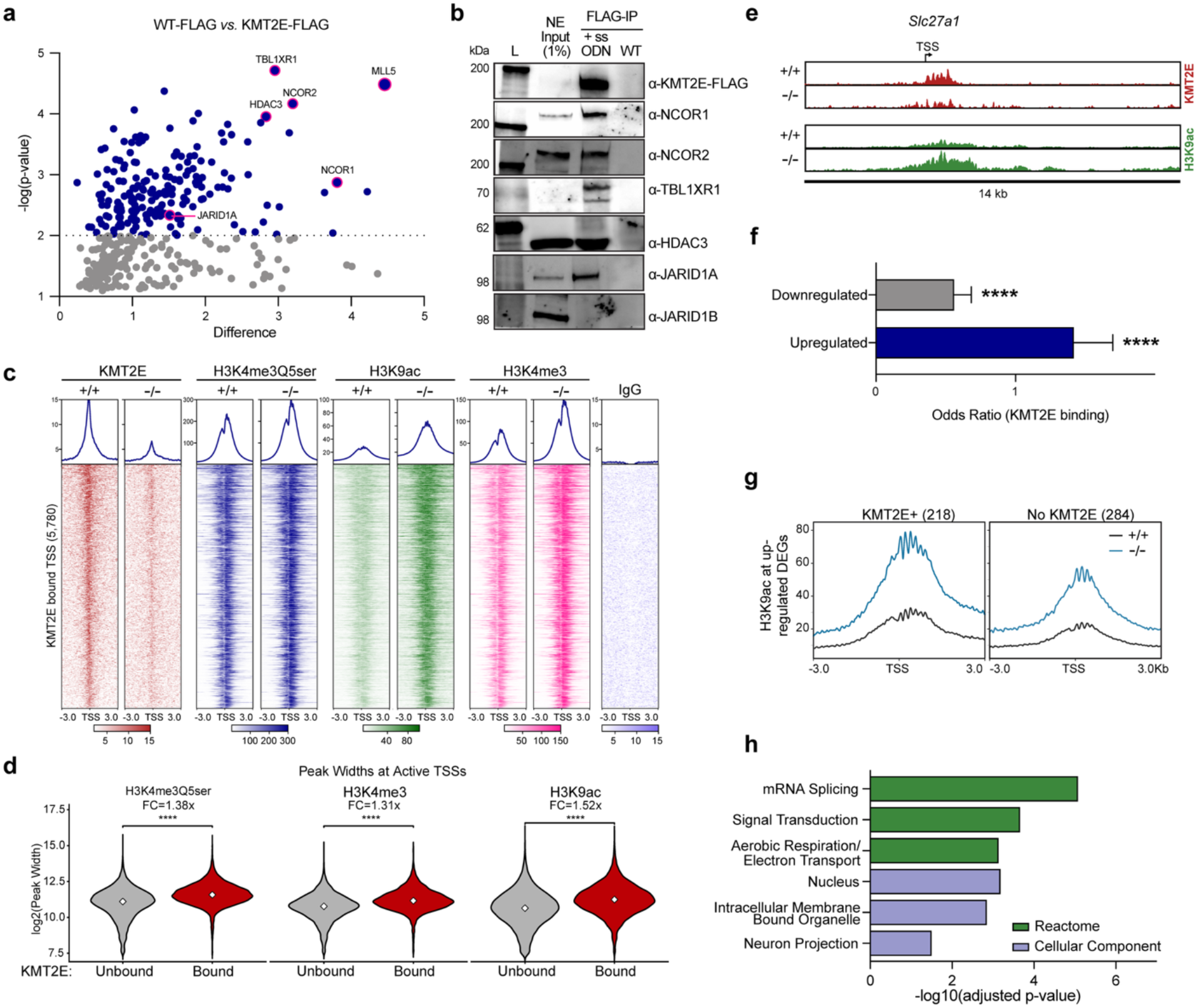
KMT2E recruits the NCoR/HDAC3 complex to restrict H3K9ac levels. **(a**) Volcano plot of KMT2E-FLAG *vs.* WT-FLAG interacting proteins from HeLa nuclear extracts identified by immunoprecipitation (IP) and LC-MS/MS. Significantly enriched proteins (*p* < 0.01, blue dots) include noted proteins of interest (labeled, pink outline). *n* = 3/group. (**b**) Immunoblotting validation confirming that KMT2E interacts with NCOR1, NCOR2, TBL1XR1, HDAC3, and JARID1A, but not JARID1B, in HeLa nuclear extracts (NE) with single-stranded oligodeoxynucleotide (ssODN) targeted to the positive (+) strand for KMT2E-3xFLAG endogenous knock-in. L, ladder. (**c**) Heatmaps and profiles for spike-in normalized CUT&RUN-seq of indicated targets (*vs.* IgG) at KMT2E-bound TSSs from P14 CB, displaying increased levels of H3K4me3Q5ser, H3K4me3 and H3K9ac in KO *vs.* WT tissues. (**d**) Violin plots showing that H3K4me3Q5ser, H3K4me3, and H3K9ac peak widths are increased at KMT2E-bound TSSs. Wilcoxon rank sum test: *p* < 2.2 x 10^-16^. Fold change (FC) of the median (KO vs. WT) is indicated for each PTM. The diamond indicates median. (**e**) Representative genome browser tracks of KMT2E and H3K9ac enrichment in WT *vs*. KO CB. For (c-e), *n* = 4/group for histone PTMs, *n* = 3/group for KMT2E. (**f**) Bar plot of odds ratio analysis examining the overlap between WT *vs.* KO DEGs at KMT2E-bound loci, demonstrating that KMT2E is enriched at upregulated DEGs. Fisher’s exact test: \*\*\*\**p* < 0.0001. Error bars represent confidence intervals. (**g**) H3K9ac levels are increased at the TSSs of upregulated DEGs in KO cerebellum, with greater enrichment observed at KMT2E-bound sites. (**h**) Significant pathways enriched for upregulated DEGs that display KMT2E enrichment at TSSs. *padj* < 0.05. Uncropped blots are provided in **Supplementary Figure 1**.

To test this hypothesis, we examined the effect of *Kmt2e* KO on these PTMs *in vivo* by performing spike-in calibrated CUT&RUN-sequencing for KMT2E, H3K4me3Q5ser, H3K9ac (a primary HDAC3 target), and H3K4me3 in WT *vs.* KO P14 cerebellar tissues; note that we focused on cerebellum in these analyses, as this region exhibited more robust transcriptional differences *vs.* mPFC. We first defined 5,780 TSSs based on KMT2E occupancy, which were marked by increased H3K4me3Q5ser, H3K9ac, and H3K4me3 in KO animals (**Fig. 3c**); such data were consistent with aberrant spreading of these marks in the absence of KMT2E-mediated repressor complex recruitment. Notably, when comparing H3K4me3Q5ser marked TSSs with KMT2E-bound (5,780) *vs.* those not bound by KMT2E (11,913), KMT2E bound loci were found to exhibit wider peak breadths at baseline for all three PTMs (**Fig. 3d, Extended Data Fig. 7a**). While both H3K4me3 and H3K9ac were found to be increased in *Kmt2e* KO animals, we subsequently chose to focus on H3K9ac as a primary mechanistic readout given the robust interactions observed between KMT2E and NCoR/HDAC3 in our mass spectrometry analyses, as well as previous findings from *Drosophila* (**Fig. 3e**).

To determine whether aberrant H3K9ac spreading underlies KMT2E-dependent transcriptional dysregulation, we next examined KMT2E occupancy at DEGs (**Extended Data Fig. 7b**). We confirmed significant enrichment of KMT2E binding at upregulated DEGs (OR > 1, **Fig. 3f**), suggesting that KMT2E likely functions to restrict expression of these genes in WT animals. At upregulated DEGs, we further observed increased spreading of H3K9ac in KO animals, as quantified by peak width (**Extended Data Fig. 7c**). In addition, H3K9ac was found to be higher at baseline and displayed more robust increases at KMT2E-bound loci (**Fig. 3g**). Notably, these effects were greater for H3K9ac *vs.* H3K4me3 or H3K4me3Q5ser (**Extended Data Fig. 7d-e**). Finally, pathway analysis of KMT2E-bound upregulated genes enriched for nuclear and neuron projection compartments, splicing, and aerobic respiration (**Fig. 3h**), directly linking *Kmt2e* loss-of-function to aberrant histone acetylation at dysregulated gene loci.

### Inhibition of H3K9ac spreading rescues transcriptional dysregulation in Kmt2e haploinsufficient neurons

Finally, we aimed to assess whether restoring H3K9ac levels may be sufficient to rescue *Kmt2e*-dependent transcriptional dysregulation. To test this in a tractable and cell-type specific system, we established primary cerebellar granule neuron cultures from *Kmt2e* WT, HET, and KO pups. These neurons were isolated from cerebellum at P7 and were differentiated for 7 days to mimic the timing of our *in vivo* analyses (**Extended Data Fig. 8a**). Consistent with our *in vivo* findings, KO neurons displayed increased H3K4me3Q5ser levels and significant transcriptional alterations, with HET neurons displaying dose-dependent epigenomic and transcriptional signatures (**Extended Data Fig. 8b-d**). Furthermore, DEGs were enriched for neurodevelopmental processes including nervous system development, synapse organization, learning, and memory, as well as pathways directly relevant to ODLURO syndrome including seizures, epilepsy, and abnormal motor learning (**Extended Data Fig. 8e**). These results confirmed that *Kmt2e* heterozygosity is sufficient to elevate histone serotonylation and dysregulate neurodevelopmental gene expression, validating our culture system as a translationally-relevant model of the ODLURO disease state.

To rescue H3K9ac levels, we targeted GCN5/KAT2A, the primary histone acetyltransferase responsible for catalyzing H3K9ac^49^. We used MB-3, a selective small molecule inhibitor^50^, with dose optimization in cultured neurons identifying 50 μM as the optimal concentration for reducing H3K9ac levels following 24-hour treatment (**Extended Data Fig. 9a**). Next, we administered 50 μM MB-3 to *Kmt2e* HET neurons, the clinically relevant genotype (**Fig. 4a**). MB-3 treatment for 24 hours significantly reduced aberrant H3K9ac spreading at KMT2E-bound loci relative to vehicle-treated HET neurons, restoring levels towards those observed in WT neurons (**Fig. 4b-c**). Transcriptional analyses further demonstrated that MB-3 treatment reversed *Kmt2e* HET gene expression patterns toward a WT-like state (**Fig. 4d, Extended Data Fig. 9b**). Examination of both up- and downregulated DEGs significantly reversed by MB-3 indicated enrichment for pathways related to nervous system development, synaptic signaling, and epilepsy (**Fig. 4e-f**). Together, these data established that rescuing aberrant H3K9ac spreading is sufficient to reverse transcriptional consequences of *Kmt2e* haploinsufficiency, additionally implicating NCoR/HDAC3-dependent histone deacetylation as a downstream effector phenomenon of KMT2E reading H3 serotonylation in developing brain (**Extended Data Fig. 9c**).

**Figure 4.**
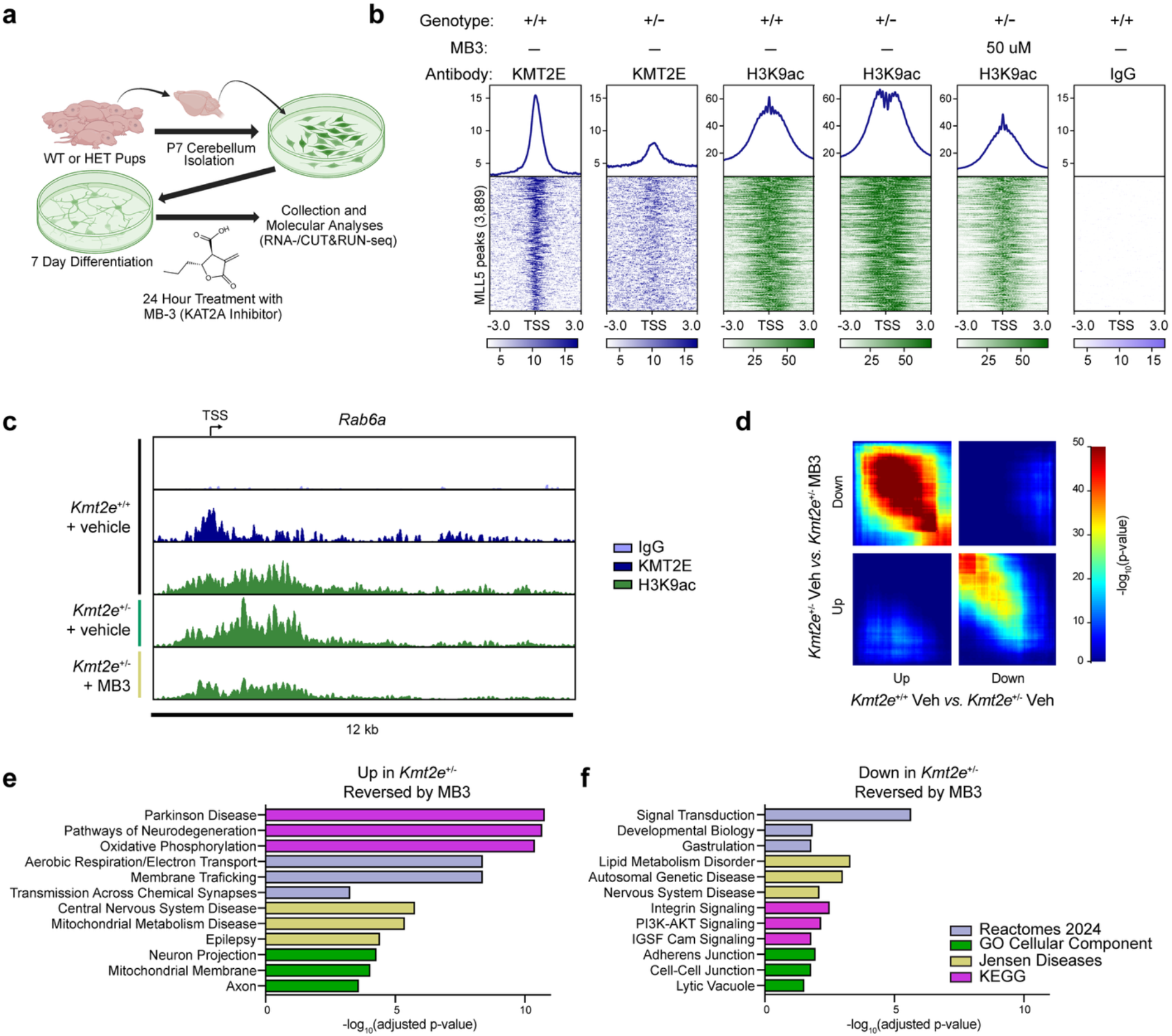
Rescue of H3K9ac levels in *Kmt2e* haploinsufficient neurons reverses transcriptional dysregulation. **(a**) Experimental workflow. Granule neurons from P7 WT or HET pups were isolated on P7, and cultured for 7 days. Neurons were treated with the KAT2A inhibitor MB-3 or vehicle for 24 hours prior to collection for RNA-seq and CUT&RUN-seq. (**b**) Heatmaps and profiles for spike-in normalized CUT&RUN-seq of KMT2E and H3K9ac (*vs*. IgG) at KMT2E-bound TSSs following vehicle or MB-3 treatment. (**c**) Representative genome browser tracks of KMT2E and H3K9ac peaks from CB granule neurons (*vs.* IgG), demonstrating that MB-3 treatment decreases H3K9ac levels. For (b-c), *n* = 3/group. (**d**) Threshold-free comparison using RRHO, showing reversal of transcriptomic patterns (upper left and bottom right quadrants) between indicated comparisons. *n* = 3/group. (**e-f**) Enrichment of significant pathways for DEGs (**e**) upregulated in HET and reversed by MB-3, and (**f**) downregulated in HET and reversed by MB-3 treatment. *padj* < 0.05.

## DISCUSSION

Our findings demonstrate the first mechanistic basis for ODLURO syndrome, revealing KMT2E as a critical *reader* of histone serotonylation that drives NCoR/HDAC3-mediated transcriptional repression at neurodevelopmental gene loci. Histone serotonylation in the developing brain matched the profiles of known active PTMs (H3K4me3 and H3K9ac), including at broad regions, which correlated with increased transcription of cell-type specific gene expression programs. In exploring *reader* proteins of histone serotonylation that may drive these gene networks, we identified the KMT2E_PHD_ domain as a putative interactor that recognizes the combinatorial H3K4me3Q5ser mark with a 5.8-fold affinity enhancement over H3K4me3 alone due to multiple distinct contacts identified via crystallography. We confirmed previous reports that the KMT2E_SET_ domain is inactive, even in the presence of its neighboring PHD domain, prompting investigations into the non-enzymatic functions of KMT2E. Previously, *in cellulo* studies using siRNA-mediated knockdown to screen for factors important for cytokinesis demonstrated that KMT2E knockdown results in similar defects to those resulting from knockdown of chromatin complex proteins NCOR2 and TBL1XR1^51^. Additionally, a recent study examining ASD proteininteraction networks identified NCoR/HDAC3 subunits enriched by KMT2E affinity purification-mass spectrometry’ from HEK-293 cells^52^. In addition to corroborating and validating the physical interaction between mammalian KMT2E and NcoR/HDAC3 as well as other chromatin-modifying proteins in these studies, our work demonstrates for the first time the biological consequence of these interactions in a relevant neuronal context. Immunoprecipitations followed by liquid chromatography-mass spectrometry from our endogenous KMT2E-tagged cell line demonstrated that KMT2E strongly interacts with the full NCoR/HDAC3 complex, as well as other epigenetic-modifying proteins (such as the H3K4 demethylase JARID1A), suggesting that it may function, at least in part, to recruit these proteins to active TSSs. Such functions appear evolutionarily conserved, as KMT2E homologues in *D. melanogaster* (*UpSET*) and *S. cerevisiae* (*SET3*/*SET4*, both of which are catalytically inactive) bind HDACs directly^26–29^. Mammalian SETD5, which similarly contains an inactive SET domain, has also been shown to recruit the NCoR/HDAC3 complex via its C-terminal intrinsically disordered region^53^. In the case of SETD5 and UpSET, their recruitment of HDAC complexes serves to keep histone acetylation in check, acting as a ‘governor’ of gene expression/activation. Similarly, our findings demonstrated that KMT2E loss-of-function perturbs H3K9ac levels and gene expression in brain during neurodevelopment. Given that aberrant spreading of histone modifications has been observed in postmortem brain tissues from patients diagnosed with autism^21^, restricting histone PTMs at key genes may be required for proficient neurodevelopment.

Our findings are also of important clinical relevance, given that *KMT2E* heterozygous mutations underlie ODLURO syndrome, a clinically severe NDD recently described in 2019^3^. Leveraging our *Kmt2e* constitutive knockout mouse model, we observed wide-ranging molecular and phenotypical alterations, including behavioral deficits that largely match the endophenotypes observed in ODLURO patients. Not only did we observe impaired cognitive outcomes, altered motor coordination, and increased seizure susceptibility, but in some instances these deficits were sex-dependent, matching the dimorphic endophenotypes observed in individuals with ODLURO syndrome. This is notable given previously reported discordance often observed between human endophenotypes and behavioral changes from genetic mouse models across monogenic NDDs^54^. In addition to previous reports indicating that *Kmt2e^-/-^* male mice are infertile^55^, we found that the proportion of surviving pups following *Kmt2e^+/-^* breeding did not conform to Mendelian genetics, strongly suggesting that the full knockout may, in some instances, lead to embryonic lethality. Consistent with this, no documented cases of biallelic mutations in the *KMT2E* locus have been reported in humans. At the molecular level, we identified robust gene expression changes in the cerebellum, a brain region associated with motor function, social interaction, and cognitive function, all of which were affected in our model. Loss of KMT2E in this region directly led to spreading of H3K9ac and H3K4me3, which associated with altered expression of key genes involved in myelination, synaptic pruning, abnormal post-synaptic currents, and chromatin remodeling pathways. We corroborated these molecular data with functional studies demonstrating altered myelination in the cerebellum, as well as altered excitatory/inhibitory currents in cerebellar granule neurons of HET and KO mice. These data support a role for KMT2E in regulating neurodevelopmental gene networks through regulation of active histone PTM spreading at histone serotonylation-marked loci. Loss of function of chromatin modifier proteins in other mouse models of ASD have shown similar synaptic plasticity and electrophysiological changes during early postnatal windows, suggesting convergent mechanisms through which disrupted chromatin regulatory programs may drive neurodevelopmental disorders.^56^

To test this mechanistic pathway within a disease context, we further showed that treatment of cultured cerebellar granule neurons with the specific histone acetyltransferase inhibitor Butyrolactone 3 (MB-3, targeting the H3K9 acetyltransferase KAT2A^57^) was sufficient to reduce H3K9ac spreading in *Kmt2e^+/-^* cells. Rescuing H3K9ac levels with MB-3 reversed discordant gene expression profiles in HET neurons, suggesting that modulation of the epigenetic landscape partially rescues the transcriptomic deficits caused by reduction of *Kmt2e* expression (**Extended Data Fig. 9c**). These data serve as proof-of-principle that inhibition of the H3K9ac machinery can alleviate the mechanistic defects caused by loss of KMT2E. Since MB-3 has a high IC_50_ (∼100 uM)^50^, systemic delivery is unlikely to achieve therapeutic concentrations *in vivo*, precluding its use in mouse models or in the clinic. Alternatively, recent engineering of targeted protein degradation tools, including the development of a selective KAT2A/KAT2B PROTAC degrader may offer a more direct therapeutic approach due to their high affinity and potency (DC_50_ ∼ 0.1-2 nM)^58^. Such outcomes will need to be rigorously tested in cells and animal models, as the technology required to deliver PROTACs across the blood-brain barrier continues to evolve^18^.

Together, our data illuminate a mechanistic axis involving histone serotonylation, KMT2E_PHD_ binding, and recruitment of epigenetic *erasers* to keep neurodevelopmental transcriptional programs intact. Constitutive disruption of this mechanistic axis severely affects development and, consequently, behavioral outcomes in mice, matching outcomes observed in human ODLURO patients. *Kmt2e* has been found to be ubiquitously expressed in all cell types in brain^59^, matching functional outcomes in both neurons and glia in the current study. Thus, further investigations into cell-type specific roles for KMT2E are needed. Additionally, patient-derived iPSC models and/or organoids may provide a critical context for understanding how patient mutations impact the mechanistic axis described here. Altogether, these findings provide fundamental insights not only into the molecular basis of ODLURO syndrome, but also broader roles for histone serotonylation and other histone modifications in NDDs.

## MATERIALS AND METHODS

### Animal housing

All mice in this study were maintained on a 12-h/12-h light/dark cycle at constant temperature (23 °C) throughout the entirety of experiments. Mice were provided with *ad libitum* access to water and food. All behavioral testing occurred during the animals’ light cycle. All animal protocols were approved by the IACUC at the Icahn School of Medicine at Mount Sinai (ISMMS) and performed in accordance with NIH guidelines. Experimenters were blinded to experimental groups in all behavioral and molecular experiments, and the order of testing was counterbalanced during all behavioral experiments.

### Generation and Validation of *Kmt2e^+/-^* Mouse Line

A synthetic single guide RNA (sgRNA; Synthego) targeting coding exon 2 of the murine *Kmt2e* locus was pre-assembled with recombinant S. pyogenes Cas9 nuclease (IDT, #1081059) to form a ribonucleoprotein (RNP) complex. The RNP complex was microinjected into C57BL/6J blastocysts to induce a double-stranded break repaired by non-homologous end-joining (NHEJ). Founder mice were screened by PCR amplification across the targeted region using primers external to the predicted cleavage site, followed by Sanger sequencing of the PCR products. Deconvolution of chromatograms was performed using ICE (Inference of CRISPR Edits)^60^. Eight primary founders carried infertile biallelic mutations, while one fertile mosaic founder (Mouse #50) was shown to harbor a 1-bp out-of-frame insertion and was therefore selected to establish the breeding line. Germline transmission of the mutant allele was confirmed by Sanger sequencing of the F1 progeny. To ensure genetic stability and minimize off-target confounders, all experimental cohorts were derived from F3 and subsequent generations. The 1-bp frameshift mutation was validated by Sanger sequencing of *Kmt2e* cDNA from an F3 homozygous KO animal. This insertion shifts the reading frame immediately 3′ of codon 60, converting the wild-type sequence (Tyr-Ala-Asp-His-Asn…) to Leu-Cys-Gly-Pro-* and introducing a premature stop codon (TAA) four codons downstream of the DSB. This mutation truncates the KMT2E protein upstream of all annotated functional domains, including the PHD finger and SET domain.

*Kmt2e* sgRNA: CTACATCGGTCTGCCCTATG

### Breeding and Genotyping

Due to observed infertility in homozygous mutant (*Kmt2e*^-/-^) male mice, the colony was maintained by pair breeding heterozygous (*Kmt2e*^+/-^) mice between 8-12 weeks of age. Litter size and date of birth were recorded for each litter, and pups were weaned at postnatal day 21 (P21). For genotyping, a 2-mm ear punch was collected by a stainless-steel ear puncher for animals aged P21 and older, whereas juvenile and adolescent mice were genotyped via toe clipping. Tissue samples were homogenized and lysed in an alkaline lysis reagent (25 mM NaOH, 0.2 mM EDTA, pH 12.0) by heating at 95°C for 1 hour. Lysates were subsequently cooled and neutralized using 1/10^th^ volume of 1M Tris-HCl (pH 8.0). To determine the genotypes, 1-2 μL of the neutralized lysate was subsequently used as template for PCR amplification. PCR was performed using Phusion High-Fidelity DNA Polymerase (ThermoFisher F530S) using primers spanning the targeted region of the *Kmt2e* locus. Amplicons were purified using a DNA clean & Concentrator kit (Zymo) and subjected to Sanger sequencing. Sequence trace files were analyzed using ICE to determine genotypes. For Mendelian distribution analysis, weanling genotypes were recorded at P21 and evaluated using a Chi-square test. For embryonic sex determination, genomic DNA (gDNA) was extracted from the tail bud of the discarded trunk tissue and subjected to PCR amplification of *Jarid1c* and *Jarid1d*.

*Kmt2e* Genotyping Fwd: AAGCTGTGCACATACTATAAATAG

*Kmt2e* Genotyping Rev: GTAATATAGCATTGTTAGGTCAAAG

*Kmt2e* Sanger sequencing primer: GACATTGTATTTGTTTTGAC

### Embryonic Tissue Collection

Adult female C57BL/6J (Jackson laboratory 664) mice were bred with age-matched males. Copulation plugs were checked every morning within 1 hour after lights on. Observation of a plug marked E0.5 and the immediate removal of the female to her own cage with a nestlet. Pregnant females were deeply anesthetized with isoflurane at the designated embryonic time point. Conceptuses were isolated from the uterine wall, and embryonic brains were enriched for the forebrain region as previously described^31^.

### Brain Tissue Collection

Adult (8-16 week) and juvenile (P14) brains were isolated, rapidly frozen on dry ice, and stored at -80°C. Whole brains were sectioned in a -20°C chamber using a 1mm coronal mouse brain matrix (Stoelting). Tissues enriched for the mPFC or cerebellum were micropunched using a 1.5 mm hollow needle (Ted Pella) according to the Allen Brain Atlas, and stored at -80°C until molecular analysis.

### qPCR

Total RNA from mouse brain tissue was extracted using the TriZol Reagent (Thermo Fisher) and cleaned up using the RNeasy Micro Kit (Qiagen). 200 ng of RNA was used to make cDNA according to the manufacturer’s protocol (iScript, Biorad 1708891), and cDNA (diluted 1:10) used for qPCR of *Kmt2e* was amplified with primers targeted to the second coding exon and normalized with *Gapdh* using the SsoAdvanced SYBR green qPCR master mix (Fisher NC1031075). Data was analyzed using the ΔΔCt method.

*Kmt2e* Fwd: CATCGGTCTGCCCTATGCG

*Kmt2e* Rev: TGCACCTGGTTACATCAGTACC

*Gapdh* Fwd: AGGTCGGTGTGAACGGATTTG

*Gapdh* Rev: TGTAGACCATGTAGTTGAGGTCA

### RNA-seq

Bulk RNA-seq assays were performed on tissues and neurons as previously described^61^. Following RNA extraction, mRNA-seq libraries were prepared using 150 ng total RNA using the Illumina Stranded mRNA Prep Kit (Illumina, #20040534) according to the manufacturer’s protocols. Library quality was measured using a Qubit Fluorometer 2.0 (Thermo Fisher) and the High Sensitivity D5000 TapeStation assay (Agilent) before sequencing on a HiSeq2500, NovaSeq 6000, or NovaSeq X system (Illumina). 25-40 million paired-end reads were targeted per sample.

### RNA-seq analysis

Raw demultiplexed fastq files were pseudoaligned and transcript abundance was quantified using Kallisto^62^ (v. 0.46.1) against the Ensembl Mus musculus reference (v. 79). Lowly expressed genes were filtered out (<10 reads across all samples), and then RUVseq^63^ (v1.32.0) were applied to account for unwanted variation among samples within each experiment that may arise from biological or technical factors unrelated to the actual experiment (such as litter, day of collection, day of library prep, etc). This uses a negative control gene set derived from the total genes identified across the sequencing experiment, after ensuring that unwanted variation did not correlate with covariates of interest, as described previously^63^. Differential expression analysis was performed using DESeq2^64^ (v1.38.3), with significant genes defined by an adjusted p-value < 0.05 and log2 fold changes where appropriate. Pairwise comparisons were performed between individual genotypes or treatments following normalization that was performed on all samples from any given sequencing experiment. For comparing across multiple genotypes, z-scores were calculated across the DEGs from a pairwise comparison of choice (i.e. WT *vs*. KO) and plotted as a heatmap with individual biological replicates in R (v4.3.0) using ggplot2^65^ (v4.0.3). For all experiments, differentially expressed genes were exported and are included in the Supplementary Tables file. Gene ontology and pathway analysis was performed using enrichR^66^ or ShinyGO^67^ (v0.81), with all protein coding genes as the background gene set. For all figures, relevant pathways were selected from the top 10 significant terms, ranked by p-value, to emphasize processes consistent with hypotheses informed by the published literature. Threshold-free gene expression overlap analysis was conducted using the Rank-Rank Hypergeometric Overlap (RRHO2) package^68^ (v1.0). Gene lists were ranked by signed p-values, calculated as the log_10_-transformed nominal p-value multiplied by the sign of the fold change. rMATS^69^ analysis was used for differential splicing analysis using default parameters.

### ChIP-seq

Chromatin immunoprecipitation (ChIP)-sequencing was performed from brain tissues and cultured neurons, as previously described.^14^ Briefly, each sample was fixed with 1% formaldehyde and rotated for 12 minutes, followed by quenching with 125 mM glycine. Samples were washed with cold 1xPBS four times, and homogenized by passing through a 22G needle at least ten times. The pellet was resuspended in cell lysis buffer (50 mM HEPES, 140 mM NaCl, 1mM EDTA, 10% glycerol, 0.5% NP-40, 0.25% Triton-X), rotated at 4°C for 15 minutes, spun down, and the supernatant removed. The pellet was then resuspended in nuclear lysis buffer (10 mM Tris pH 8, 200 mM NaCl, 10 mM EDTA, and 0.5 mM EGTA), rotated at 4°C for 10 minutes, spun down, and the supernatant removed. The final pellet was resuspended (10 mM Tris pH 8, 100 mM NaCl, 10 mM EDTA, 0.5 mM EGTA, 0.1% sodium deoxycholate, 0.5% N-lauroylsarcosine) and sonicated using a Covaris E220 for 30-60 minutes at 4°C with the following conditions: peak incident power, 140; duty factor, 10%; cycles/burst, 200; water level, 0. Triton-X was added (final concentration 1%), and equal amounts of chromatin were incubated with 2.5 μg of H3K4me3Q5ser antibody (MilliporeSigma ABE2580) bound to protein A Dynabeads (ThermoFisher 10001D) at 4°C overnight. Samples were then washed 8 times with RIPA buffer and once with TE Buffer with 50mM NaCl. The DNA was eluted (50 mM Tris pH 8, 1% SDS, 10 mM EDTA) and reverse crosslinked at 65°C. RNA and protein was digested, and DNA purified using Zymo ChIP clean and concentrator. 1% input was removed prior to antibody addition and purified in parallel as a control. Libraries were generated using the TruSeq ChIP library preparation kit (Illumina IP-202-1012) according to manufacturer’s protocol. Quality control of libraries was assessed with the High Sensitivity D5000 TapeStation assay (Agilent), prior to sequencing on an Illumina HiSeq2500 or NovaSeq6000.

### ChIP-seq analysis

Raw fastq files were aligned to the mm10 mouse genome using the NGS Data Charmer pipeline with default settings (HISAT v.0.1.6b). Peak calling was performed using MACS2^70^ (v.2.1.0) with --broad (q < 0.1), and filtered for peaks with FDR < 0.05. All peaks were annotated using the mm10 genome with the Homer^71^ package (v4.10). Functional annotation analysis of uniquely annotated loci was conducted using ShinyGO^67^ (v0.81) with a background of all protein-coding genes in the mm10 genome. Visualization of differential peaks were accomplished using deepTools^72^ (v3.5.3).

### CUT&RUN-seq

Cleavage Under Target & Release Under Nuclease (CUT&RUN)-sequencing was performed from cultured neurons and brain tissues, as previously described^10^. For cultured neurons, cells were scraped off plates in ice cold PBS, pelleted by centrifugation at 4°C and 1100 x g for 5 min prior to isolating nuclei by resuspending in 500 μL of nuclear extract buffer (NEB: 20 mM HEPES-KOH, pH 7.9, 10 mM KCl, 0.5 mM spermidine, 0.1% Triton X-100, 20% glycerol and freshly added protease inhibitors (Halt Protease Inhibitor Cocktail, EDTA-free, Thermo Fisher Scientific) thoroughly. For brain tissues, tissue punches were homogenized using plastic pestles (Fisher, 12-141-364) in 200 μL of NEB, volume increased to 500 μL, and then passed through a 21 gauge needle 10 times. For both, nuclei were pelleted and resuspended in 500 μL of NEB and counted using a hemocytometer, and 50k (histone modifications) or 100k (KMT2E) nuclei were used per biological replicate.

Nuclei were bound to activated Concanavalin A beads (Polysciences, 86057-3) for 10 minutes at room temperature, washed (WB: 20 mM HEPES, pH 7.5, 150 mM NaCl, 0.1% Triton X-100, 0.1% Tween-20, 0.5 mM spermidine, 0.1% BSA, freshly added protease inhibitors) 2 times, and primary antibody (see table below) added for overnight incubation in antibody buffer (WB with 2 mM EDTA) on a mixer (tubes on their side at ∼20 degree upward angle). The second day, nuclei were washed and resuspended in 50 μl of cold wash buffer, and 2.5 μl of pAG-MNase (Epicypher, 15-1016) was added and incubated for 1 h at 4 °C on the same mixer. Nuclei were then washed two times with 1 ml ice-cold wash buffer, followed by one wash in 1 ml low-salt rinse buffer (20 mM HEPES, pH 7.5, 0.5 mM spermidine, 0.1% Tween-20 and 0.1% Triton X-100). Nuclei were resuspended in ice-cold calcium incubation buffer (3.5 mM HEPES, pH 7.5, 10 mM CaCl_2_, 0.1% Tween-20, 0.1% Triton X-100) on ice, and incubated for 30 min, with the reaction being quenched by adding 100 μl of 2× stop buffer (340 mM NaCl, 20 mM EDTA, 5 mM EGTA, 0.1% Tween-20, 0.1% Triton X-100, 25 μg ml−1 RNase A (Thermo Fisher Scientific) and 0.1 ng per 100 μl of E. coli spike-in DNA (Epicypher, 18-1401)) Nuclei were incubated at 37 °C for 15 min with no shaking to allow for release of chromatin and digestion of RNA. Beads were placed onto a magnet, and the supernatant (200 μl) was collected. DNA was isolated using the Zymo ChIP DNA Clean & Concentrator kit (D5205) and eluted in 30 μl and frozen at −20 °C for library preparation.

Library preparation was performed using the NEBnext Ultra II DNA library kit (E7645L) with multiplexed adapters with minor modifications. CUT&RUN DNA underwent end repair and adapter ligation according to the manufacturer’s protocol (1:15 adapter dilution was used). DNA was amplified using 16 PCR cycles with 10 s of extension time per cycle. Libraries were quantified using the Qubit fluorometer (Thermo Fisher Scientific), and the library size distribution was checked using the Tapestation DNA High Sensitivity ScreenTape (Agilent). Libraries were pooled at an equimolar concentration and sequenced on the Illumina NovaSeq 6000 or X+ sequencer by the NYU Genome Technology Center, with 20-30 million reads targeted.

### CUT&RUN-seq analysis

Raw Fastq files were aligned to the mm10 genome using bowtie2^73^ (v2.5.0). Low-quality reads were filtered using Samtools^74^ (v.1.9) with a MAPQ cut-off score of 30, and only unique, deduplicated reads were retained for further processing. Bigwig files were produced using deepTools^72^ (v.3.5.1), using an ENCODE^75^ mm10 v2 blacklist file to discard the regions with consistently non-specific signal. Bigwigs were scaled using *E. coli* spike-in controls to normalize sequencing depth. To determine normalization factors based on *E. coli* reads, each sample was aligned to the *E. coli* genome (MG1655), and the unique deduplicated reads were compared across genotypes and/or treatment conditions, per antibody, per experiment. The sample with the lowest number of *E. coli* reads was determined, and all samples were scaled by dividing their corresponding *E. coli* read count by this minimum number. For each group, the bam files were merged and peak callling was conducted using MACS2^70^ (v2.1.0) with the corresponding merged IgG file as a control, and filtered for peaks with FDR < 0.05 or greater where appropriate. For histone modifications, broad mode was used for peak calling. For KMT2E, both broad and narrow mode were used, and then subsequently merged using bedtools^76^ merge (-d 501). Gene annotation was conducted by overlapping peaks with all known transcript start sites using bedtools intersect (2.31.0), using the mouse mm10 Ensembl GENCODE vM23 list. Heatmaps were made using the deepTools (v3.5.5) packages. For deepTools, heatmaps were anchored to the notated peak list, either focusing on all KMT2E -bound TSSs, all H3K4me3Q5ser+ TSSs, or DEG overlapping TSSs. ChEA analysis on annotated peaks was conducted using EnrichR^66^ with a significance threshold of adjusted p < 0.05.

### Comparison of Peak Widths

In order to calculate peak widths, a custom enrichment analysis was performed. Deeptools^72^ multiBigwigSummary was run for the histone PTM-seq and matching input files of choice, using the bins option. Binsize was set to 500, with a sliding window of -n set to -450, to allow for sliding windows of 500 basepairs in size at every 50 basepairs across the entire mm10 genome (excluding blacklisted regions). Then, the raw counts .tab file was further analyzed in R (4.5.0). The average and standard deviation of each bigwigs’s bins was calculated, and cutoffs used to determine which bins to keep as enriched in K9ac signal. For H3K9ac peak width from P14 cerebellum, the minimum signal over a bin was set as one standard deviation above the mean for the WT H3K9ac signal. Additionally, the minimum input signal was set as 2 standard deviations below the average, to rule out regions with little to no signal. Finally, a ratio in signal between the histone PTM and corresponding input, and a minimum fold change enrichment of 2 fold was required. Following identification of these enriched bins, overlapping and neighboring bins were merged together using bedtools merge (-d 501, to join nearby bins together). The final .bed file was sorted and inspected in Integrative Genomics Viewer (IGV) to ensure accurate peak calling at both the top and bottom 50 peaks. The cutoffs used for the WT, input, and ratio were kept consistent across the peak width calling for the other genotypes/conditions in order to capture regions where peak widths ‘spread.’ Peaks were merged with TSSs using bedtools intersect (2.31.0), using the mouse mm10 Ensembl GENCODE vM23 list, and subsequently separated out based on select criteria (i.e. KMT2E bound vs. unbound). For quantitative comparison of the peak widths, statistics were calculated between groups using dplyr^77^, and boxplots generated using ggplot2^65^ (4.0.3). Statistics between groups were calculated using the Wilcoxon rank-sum test, assuming unpaired samples and a continuous distribution.

### Production of modified histone peptides

Modified peptides corresponding to H3_1–10_ were prepared, as previously described^1^. Briefly, peptides were synthesized on 2-chlorotrityl resin using manual addition of the reagents (using a stream of dry N2 to agitate the reaction mixture), incorporating a C-terminal Lys(biotin) residue. The resin was swollen using 5 mmol of DIEA in DCM (1 x 1 hour). The coupling of the C-terminal Lys(biotin) residue was performed utilizing 1 eq. of Fmoc-Lys-Biotin and 5 eq. of DIEA in DCM (1 x 4 hour). Following coupling, a capping step utilizing 17:2:1 (DCM:MeOH:DIEA, 5 mL) was performed (1 x 10 min). For manual solid phase synthesis, typical cycles were: (i) FMOC group deprotection with 3 mL of 20% piperidine in DMF (1 × 1 min, 2 × 8 min) and (ii) coupling of 5 eq. amino acid to the growing peptide chain with 4.9 eq. of HOBt/HBTU and 10 eq. DIPEA for 45 min. Serotonylated glutamine residues were introduced by incorporation of a Fmoc-L-glutamic acid γ-allyl ester (Chem-Impex International). The allylic ester was deprotected utilizing Pd(PPh3)4 (0.25 eq) and dimethyl barbituric acid (5 eq) in 5 mL of DCM. This mixture was agitated for 30 minutes, washed, and repeated. The resin was washed to remove remaining catalyst using DMF, 500 mg of Cupral in 50 mL DMF, 1% DIEA in DMF (50 mL), 0.1 M HOBT in DMF (50 mL), DMF, and DCM. Serotonin was coupled by treating resin (25 µmol) with PyAop (1.2 eq) and DIEA (5 eq) in DMF (1 mL) for 2 minutes. To this was added serotonin (2 eq) and DIEA (5 eq) in DMF (1 mL). This was agitated with N2 for 30 minutes. Upon completion of serotonin coupling, a cleavage was performed using 92.5% TFA, 2.5% EDT, 2.5% TIS and 2.5% H2O. The crude peptide was then precipitated with diethyl ether and allowed to air dry. Peptides were resuspended in diH_2_O, aliquoted, and frozen at -80 until use.

### Methyl Reader Array

A library of recombinant methyl *reader* domains was generated and used, as previously described^33,78^. This version of the array harbors a collection of different GST fusion proteins, including fusions to Tudor domains, Chromo domains and PHDs. Briefly, *reader* domains were cloned into pGEX vectors as fusion proteins with glutathione-S-transferase (GST), followed by expression and purification from *E. coli*. An array of purified domains was robotically arrayed onto nitrocellulose-coated glass slides (Grace Biol-labs, Cat #305170), with each grid containing 12 fusion proteins (200 ng of each) arrayed in duplicate, with a GST only control. C-terminal biotinylated H3_1-10_ peptides with corresponding modifications (H3unmod, H3K4me3, H3Q5ser, H3K4me3Q5ser) were pre-conjugated using FluoroLink (Cy3), hybridized to the microarray (pre-blocked), and residual peptides washed off. Finally, the slide was scanned using a microarray analyzer (Genomic Solutions).

### Cloning and Protein Purification

Human cDNA from HeLa cells was utilized to clone *KMT2E* PHD (114-166), SET (323–472), and PHD-SET (107–473) into the pET28a backbone using Gibson cloning and HiFi DNA assembly master mix (NEB E2621). All constructs were verified by Sanger sequencing, and transformed into *Escherichia coli* BL21 cells for expression. Protein expression was induced at an OD-600 of 0.6-1.0 with 0.25 mM isopropyl β-D-1-thiogalactopyranoside (IPTG) at 16°C overnight in LB medium. Cells were harvested and resuspended in the lysis buffer containing: 500 mM NaCl, 20 mM Tris-HCl (pH 7.5), 0.1% Triton-X 100, 5 % glycerol, 1 mM DTT, and protease inhibitors (1X Halt Protease Inhibitor, Thermo 78430). Cells were lysed and clarified by centrifugation, and supernatant was applied to a HisTrap affinity column (GE Healthcare). The column was then washed with five column volumes of lysis buffer before bound proteins were eluted with buffer containing: 150 mM NaCl, 20 mM Tris-HCl (pH 7.5), 2.5 mM DTT, 5% glycerol, 0.1 % Triton-X 100, and 300 mM imidazole. Purified proteins were buffer-exchanged into storage buffer (150 mM NaCl, 20 mM Tris-HCl, pH 7.5, 1 mM DTT, 2.5 % glycerol, 0.05% Triton-X 100), concentrated to approximately 5-10 mg mL⁻¹, aliquoted, and stored at −80 °C for future use. For crystallization and isothermal titration calorimetry, purified proteins underwent further purification using additional methods. Eluted protein was purified by anion-exchange chromatography using a HiTrap™ Q HP column (Cytiva) and size-exclusion chromatography on a Superdex 75 column (Cytiva). H3 peptides for isothermal calorimetry, reader array and pulldown assays were prepared as described above or synthesized commercially (Beijing SciLight Biotechnology).

### Isothermal Titration Calorimetry

Isothermal titration calorimetry (ITC) experiments were performed at 25°C using the MicroCal PEAQ-ITC instrument (Malvern Instruments). The sample cell was loaded with 200 μL of KMT2E-PHD protein (100 μM), and titrated with 17 sequential injections of histone peptide (H31_-15_) with the respective modification at a concentration of 1 mM. Titration data were analyzed using Origin 7.0 software and fitted to a single-site binding model. ITC statistics are provided in **Extended Data Table 1.**

### Crystallization, data collection, and structure determination

Crystallization trials were performed using the sitting-drop vapor diffusion method at 18°C. Purified *KMT2E-*PHD protein (20 mg mL⁻¹) was preincubated with H3K4me3Q5ser_1-7_ peptides at a 1:5 molar ratio in buffer containing: 100 mM NaCl and 20 mM Tris-HCl (pH 7.5). Crystals of the KMT2E-PHD–H3K4me3Q5ser complex were obtained under multiple reservoir conditions, including mixtures containing carboxylic acids, Bicine/TRIS buffers, hexylene glycol, and polyethylene glycol (PEG) 1000 and PEG 3350 at pH 8.5.

Crystals were cryoprotected and flash-frozen in liquid nitrogen prior to data collection. X-ray diffraction data were collected at the Shanghai Synchrotron Radiation Facility. Diffraction images were indexed, integrated, and scaled using HKL2000^79^. Structures were solved by molecular replacement using MOLREP^80^ from the CCP4 suite, with the KMT2E-PHD finger in complex with H3K4me3 structure (PDB ID: <u>4L58</u>) as the search model. Model building and refinement were carried out using COOT^81^ and PHENIX^82^, respectively. Data collection and refinement statistics are provided in **Extended Data Table 2**.

### Histone Peptide Pulldowns

Histone tail peptide pulldowns were performed from nuclear extracts and using recombinant proteins, as previously described^10^. Briefly, biotinylated unmodified H3, H3Q5ser, H3K4me3, or H3K4me3Q5ser peptides (2 ug; 1-10 length) were mixed with 20 μL of prewashed M280 streptavidin beads (ThermoFisher 11206D) in 0.1% PBS/Triton-X 100 for 1 hour rotating at RT. For nuclear extract peptide pulldowns, 1 mg of soluble nuclear extract was pre-cleared with non-conjugated beads for 30 minutes at 4°C, then incubated overnight at 4°C with rotation with peptide-conjugated beads. The following day, beads were washed 6X in medium salt conditions (300 mM NaCl) and proteins eluted and western blotting performed as described below to assess the amount of KMT2E bound. For recombinant KMT2E-PHD-SET protein, 4 ug of purified HIS-tagged protein was added and incubated for 2-3 hours at 4°C (rotating). Samples were washed with medium salt conditions (300 mM NaCl) 6X and eluted, and western blotting performed. Quantification of western blotting was performed using ImageJ.

### Western Blotting

All samples for western blotting were mixed to a 1X final concentration with reducing SDS sample buffer (ThermoFisher J61337.AD) and boiled for 8 minutes. Input and pulldown samples were run on 4-12% Bis-Tris gels (Invitrogen, NW04122) for 45 minutes at 150V. Proteins were then transferred to 0.2 µm nitrocellulose membranes using a Trans-Blot Turbo Transfer System (BioRad) according to the manufacturer’s protocol. The membrane was blocked in 5% milk (TBS) for 1 hour, and mixed with primary antibody (in 1.5% milk in TBS, see table below) overnight at 4°C. Blot were washed 3X for 15 minutes in TBS-T, and then incubated with secondary antibody (see below table) at room temperature for 1 hour, followed by washing 3X for 15 minutes in TBS-T. Blot were imaged using a BioRad ChemiDoc MP, and quantification of western blotting was performed using ImageJ.

### Behavioral Phenotyping

All animals (8-30 weeks) were handled for a minimum of 2 minutes on two consecutive days before initial behavioral testing. Animals were habituated to the testing room for 1 hour before each behavioral assay. All testing occurred during the light phase and at consistent times of day across multi-day paradigms to minimize confounding variables. All behavioral experiments were run in cohorts that included all genotypes to ensure reproducibility and rigor, with a maximum of 2 genotypes per sex per litter to control for litter effects. Researchers were blinded to genotype until data collection and analysis were complete. Behavioral data are presented by sex when sex effects were statistically detected, and by genotype (collapsed across sex) when no sex differences were observed.

### Statistics

Statistical analyses for behavioral and immunoassay data were conducted using Prism software (GraphPad, v.10.4.1). Data distributions were assessed for normality, and normally distributed data were analyzed using parametric tests, while non-normally distributed data were analyzed using non-parametric alternatives. For experiments involving multiple conditions, one-way or two-way ANOVAs were performed, including *post hoc* analyses when appropriate. For time course analyses with multiple measurements from the same animal, repeated measure ANOVAs were performed. Two-tailed Student’s t-tests were used for comparisons between two conditions. Grubb’s test (alpha = 0.05) was applied to detect outliers where necessary. Statistical significance was defined as p ≤ 0.05. For genomics analyses, statistical cutoffs were used as indicated throughout. For comparing large datasets, such as peak intensity and peak width, Wilcoxon rank sum tests were performed and p-values calculated.

### Open Field

Mice were placed into a 16×16 square arena with dim lighting for 5 minutes. The total distance and amount of time spent in the center *vs.* periphery were automatically recorded by Fusion software.

### Light Dark Box

Anxiety-like behavior was tested using an apparatus containing two interconnected 20×20 cm compartments (Omnitech Electronics Inc.). One compartment was illuminated for the session (“light” side), while the other was blocked by an opaque black perspex lid (“dark” side). The total distance, time spent, and number of crossovers between each compartment were automatically recorded by Fusion software over a 10-minute testing duration.

### Y-Maze

Mice were placed in one arm of a standard Y-maze and allowed to explore freely for 8 minutes. Sessions were recorded via overhead camera, and arm entry sequences were scored manually. Spontaneous alternation was calculated as the number of triads containing entries into all three arms divided by the total number of possible triads (total arm entries − 2), expressed as a percentage.

### 3-Chamber Test

Social behavior was assessed using a three-chambered sociability apparatus with acrylic walls and a white matte bottom (Noldus). On the day of testing, subjects were first habituated to the empty apparatus for 10 minutes, followed by the apparatus containing two empty wire corrals for 10 minutes. During the test phase, an age-, sex-, and strain-matched stranger mouse was placed under a wire corral in one chamber and a novel object was placed under a wire corral in the opposite chamber. Subject mice were allowed to freely investigate for 10 minutes. Behavior was recorded and analyzed using EthoVision XT (Noldus, v9.14). Social and object interaction times were defined as the duration during which the subject’s nose point was detected within a circular interaction zone surrounding the respective corral.

### Auditory Fear Conditioning

On day 1, mice were habituated to the conditioning chamber, consisting of a stainless steel grid scented with 70% ethanol, for 10 minutes. On day 2, following a 3-minute baseline measurement, mice underwent auditory fear conditioning consisting of six tone-shock pairings in the same chamber. Each 20-second tone (2 kHz, 80 dB) co-terminated with a 2.0-second, 0.7 mA foot shock, with an inter-trial interval of 60 seconds. Over the subsequent two days, extinction was conducted using 20 tone-alone presentations per day with an inter-trial interval of 40 seconds. The first day of extinction occurred in the conditioning context, while the second day occurred in an alternate context consisting of a curved plastic insert and a plastic floor scented with 1% Micro-90. On the final day, fear recall was assessed with four tone presentations in the alternate context.

### Rotarod

Motor coordination and motor learning were assessed using an accelerating rotarod paradigm. Mice were placed on the rotarod apparatus and subjected to an acceleration protocol beginning at 4 rpm for 2 minutes, followed by a linear acceleration to 80 rpm over a maximum trial duration of 10 minutes. Latency to fall was recorded for each trial. Testing was conducted over 4 consecutive days with 3 trials per day, and an inter-trial interval of at least 15 minutes was maintained to minimize fatigue effects. The average latency to fall across the three daily trials was used for analysis.

### Kainic Acid-Induced Seizure Susceptibility

Seizure susceptibility was assessed using an escalating dose kainic acid (KA) paradigm as previously described^83^. All experimenters were blinded to genotype throughout the experiment. Mice were weighed, tail-marked for identification, and placed individually into clean cages without bedding. Kainic acid (2 mg/mL in sterile saline) was administered intraperitoneally at an initial dose of 10 mg/kg, and the time of injection was recorded. Mice were observed for convulsive seizure activity for 10 minutes following each injection. Seizure behavior was scored using the Racine scale. If no convulsive seizure (Racine scale ≥ 3) was observed within the 10 minute window, a booster dose of 5 mg/kg was administered and observation continued for an additional 10 minutes. This procedure was repeated until either a Racine scale ≥ 3 seizure was observed or a cumulative dose of 40 mg/kg had been administered. Upon observation of a convulsive seizure (Racine scale ≥ 3), the booster schedule was discontinued. Mice were monitored for an additional 60 minutes following the final injection, during which the timing and severity of each seizure event were recorded. Mortality was noted where applicable. Mice were euthanized at the conclusion of the monitoring period.

### Immunofluorescence and analysis

Male P14 mPFC sections were cut at 40 μm, permeabilized in 0.3% PBST for 30 min, and then blocked in 10% normal donkey serum for 2 h at room temperature. Sections were incubated overnight at 4°C with 1:250 anti-Tbr1 (Abcam, ab183032), or co-stained with 1:100 anti-Brn1 (Novus Biologicals, NBP1-49872) and 1:100 anti-Ctip2 (Abcam, ab28448). On the following day, sections were washed three times in 1× PBS for 10 min each and incubated for 2 h at room temperature with 1:1000 secondary antibodies, including donkey anti-goat Alexa Fluor 647 for Brn1, donkey anti-rabbit Alexa Fluor 568 for Ctip2, and donkey anti-rabbit Alexa Fluor 546 for Tbr1 (Thermo Fisher). Sections were then washed, counterstained with 1:10,000 DAPI, and mounted with ProLong Gold Antifade Mountant (Thermo Fisher P36934). Images were acquired on a Zeiss LSM 780 upright microscope using a 40x objective with Zen Black software, with 3×4 tiled images. For cell counting, z-stacks were saved in .tiff format and analyzed using ImageJ/FIJI software. Images were thresholded using the Otsu method. Binary masks were generated, holes were filled, and watershed segmentation was applied where necessary to separate clustered nuclei. Regions of interest were selected manually and applied across matched images using the ROI Manager. Within the selected region, particles were quantified using Analyze Particles with a size threshold of 40–infinity µm^2^ and circularity thresholds of 0-1 for Brn1 and Tbr1, and 0-0.3 for Ctip2. All image processing and quantification parameters were applied identically across experimental groups.

### Patch-Clamp Electrophysiology

*Kmt2e^-/-^* mice (3-6 weeks old) were deeply anesthetized with isoflurane and decapitated. Brains were rapidly removed and chilled in cutting artificial CSF (ACSF) containing (in mM): N-methyl-D-glucamine 93, HCl 93, KCl 2.5, NaH2PO4 1.2, NaHCO3 30, HEPES 20, glucose 25, sodium ascorbate 5, thiourea 2, sodium pyruvate 3, MgSO4 10, and CaCl2 0.5, pH 7.4. Brains were embedded in 2% agarose and coronal slices (300 µm thick) were made using a Compresstome (Precisionary Instruments). Brain slices were allowed to recover at 31 ±1 °C in cutting solution for 30 minutes and thereafter at room temperature in holding ACSF, containing (in mM) : NaCl 92, KCl 2.5, NaH2PO4 1.2, NaHCO3 30, HEPES 20, glucose 25, sodium ascorbate 5, thiourea 2, sodium pyruvate 3, MgSO4, and CaCl2 2, pH 7.4. After at least 1 hour of recovery, the slices were transferred to a submersion recording chamber and continuously perfused (2 -4 ml/min) with ACSF containing (in mM) : NaCl 124, KCl 2.5, NaH2PO4 1.2, NaHCO3 24, HEPES 5, glucose 12.5, MgSO4 2, and CaCl2 2, pH 7.4. 10µM CNQX + 1µM TTX or 100μM Gabazine + 1µM TTX were added at least 30 minute before mIPSC or mEPSC recordings, respectively. All solutions were continuously bubbled with 95% O2/5% CO2. Cerebellar granule cells were visually identified with infrared differential contrast optics (BX51; Olympus). Whole-cell patch clamp recordings were performed at room temperature using a Multiclamp 700 A amplifier (Molecular Devices). Recording electrodes (5-7 MΩ) pulled from borosilicate glass were filled with solution containing (in mM): Cs gluconate 122, HEPES 10, KCl 5, MgATP 5, Na2GTP 0.5, QX314 1, and EGTA 1, pH 7.25. Data acquisition (filtered at 10 kHz and digitized at 10 kHz) and analysis were performed with pClamp 11 software (Molecular Devices). mEPSCs and mIPSCs were recorded in voltage clamp mode at a holding potential of -80mV and +10 mV, respectively, for 3 minutes following breakthrough. Only cells with stable input resistances were included in the analysis. Recording and analysis were conducted in blinded manner.

### Electron Microscopy Sample Preparation and Imaging

For electron microscopy, animals were sacrificed by transcardial perfusion with 1% PFA flush for 1 minute at 3-5 ml/minute, then switched to Karnovsky’s fixative (2% paraformaldehyde and 2% glutaraldehyde in 0.1M sodium cacodylate) at 5 mL/min for 10 minutes. Brains were then resected and post-fixed in heavy fixative for at least 1 week. Brains were sectioned at 200 µm on a vibratome, and sections were stored in Karnovsky’s fixative until further processing. Samples were processed for transmission electron microscopy (TEM) using an enhanced heavy metal staining and Epon embedding protocol. Tissues were rinsed in 0.1 M sodium cacodylate buffer and incubated in 0.1% tannic acid for 30 min at room temperature, followed by washing and post fixation in 2% osmium tetroxide/1.5% potassium ferricyanide in cacodylate buffer for 30 minutes in the dark. After rinsing in double-distilled water, samples were sequentially treated with 1% thiocarbohydrazide and 2% osmium tetroxide (30 minutes each, room temperature, dark), with intervening washes. En bloc staining was performed with 1% aqueous uranyl acetate (1–2.5 h, room temperature in darkness), followed by incubation in lead aspartate (30 minutes, 60°C). Samples were washed extensively in water and dehydrated through a graded ethanol series, transitioned through propylene oxide, and infiltrated with EMbed-812 resin using increasing resin:propylene oxide ratios. Specimens were embedded in fresh resin and polymerized at 60– 65°C for 48–72 hours. Ultrathin sections (70–90 nm) were collected on copper grids, stained with uranyl acetate and lead citrate, and air-dried prior to imaging. Samples were imaged using a JEOL JEM-1400 TEM at 75 kV, and images were collected using a NANOSPRT12 digital camera. 8 low-resolution (800× electron micrograph) images were analyzed by quantifying the number of myelinated axons per visual field. 30 high-resolution (5000x electron micrograph) images was quantified per animal (a minimum of 53 axons/animal) and MyelTracer was used to calculate g-ratio^84^.

### Methyltransferase Assays

Methyltransferase activities of recombinant KMT2E_PHD_, KMT2E_SET_, and KMT2E_PHD+SET_ were measured using the MTase-Glo Methyltransferase Assay (Promega, V7601), according to the manufacturers protocol, with the MLL4 complex used as a positive control. The assay detects S-adenosyl-L-homocysteine (SAH) produced during methyl group transfer through a coupled bioluminescent reaction. Reactions were assembled in a white, 96 well plate in a final volume of 20 µL containing 20 µM SAM and 1 µg of histone peptide (H3 1–10, unmodified) in the MTase-Glo reaction buffer. Each protein was titrated using two-fold serial dilution across a final concentration range of 1.3 to 170 nM, with a no-enzyme negative control included on every plate. Plates were sealed and incubated at 37 °C for 1 hour, after which MTase-Glo Reagent and Detection Solution (5 μL) were added and mixed, and kept at 23°C for an additional 30 minutes. Luminescence was recorded on a SpectraMax iD5 plate reader with an integration time of 1 s per well. Background luminescence from no-enzyme controls was subtracted, and the signal was plotted against protein concentration. For reactions used in western blotting analysis with H3K4me specific antibodies, a 20 μL reaction was set up using 1 ug of recombinant histone H3.1 (Active Motif 31294) and 100 nM of the indicated protein. The reactions were incubated for 1 hour at 37°C, and quenched by resuspending in 1X lamelli reducing load dye and boiled for 5 minutes. The western blot was performed as described above, with the H3K4me1/H3K4me2/H3K4me3 primary antibodies mixed (1:1000 for each, see antibody table).

### Generation of KMT2E-3XFLAG HeLa Cells

Endogenous knock-in of a 3X FLAG tag on the C-terminus of the endogenous *KMT2E* locus was planned using the Invitrogen TrueDesign Genome Editor online tool. Guide RNAs and ssODNs targeting the C-terminus were ordered from Integrative DNA Technologies. A ribonucleoprotein (RNP) complex was assembled by combining 1.2 µL sgRNA (120 pmol) with 1.64 µL recombinant S. pyogenes Cas9 nuclease (IDT Alt-R Cas9, 62 µM stock; 100 pmol final) in PBS to a final volume of 5 µL, mixing gently by pipetting, and incubating for 10–20 min at room temperature. A 100-mer to 200-mer single-stranded oligodeoxynucleotide (ssODN) donor encoding the desired edit was resuspended in 1× TE to 100 µM, and 1 µL (100 pmol) was added to the nucleofection mix immediately prior to electroporation, along with 1 µL of 100 µM Alt-R electroporation enhancer (4 µM final). HeLa-S3 cells (ATCC CCL-2.2) at 40-70% confluency were harvested by trypsinization, washed once in PBS, counted, and pelleted at 500 × g in a swinging-bucket rotor. Two hundred thousand cells per reaction were resuspended in 20 µL nucleofector solution (Lonza), combined with the pre-assembled 5 µL RNP mix (and 1 µL ssODN + 1 µL enhancer for HDR reactions; total volume 27 µL), and transferred to a Lonza 16-well Nucleocuvette Strip, avoiding bubble formation. Nucleofection was performed using the cell line– optimized program (SE-DS150). Immediately following nucleofection, 75 µL of pre-warmed culture medium was added to each well, and cells were transferred to a 6-well plate containing pre-warmed medium. For HDR reactions, the recovery medium was supplemented with 1 µM IDT Alt-R HDR Enhancer V2 (diluted 1:690 from a 690 µM stock); medium was replaced with standard culture medium 18-24 hours post-nucleofection. Mock-electroporated cells and a parallel GFP-transfected control (1 µL GFP plasmid provided with the Lonza kit) were included in each experiment to monitor viability and nucleofection efficiency. Following expansion of the cells (72-96 hours), a small aliquot was collected for genotyping. gDNA was isolated and PCR performed using primers spanning the targeted region, and PCR products run on a 2 % agarose gel. Sanger sequencing of the PCR products was performed to confirm correct sequence, and western blotting performed for FLAG. All PCRs, Sanger sequencing, and western blots were compared to WT cells which were mock nucleofected with GFP only.

CRISPR-Cas9 Guide RNA:

ACAATTACCATGGGTCAGGG

PAM=TGG, Score 90.17, edit site distance 9 bp).

ssODN- (colored=homology arms):

AGAACATTTAAAAAAATGTTTTTGGAGTCCATTTTAATGCTTGTCATCGTCGTCTTT GTAGTCGATGTCGTGATCCTTATAATCGCCGTCGTGGTCCTTGTAGTCCCACCCTGACCCATGGTAATTGTTTTGAAATGTTGGTGGCACCTGT

ssODN+ (colored=homology arms):

CAGGTGCCACCAACATTTCAAAACAATTACCATGGGTCAGGGTGGGACTACAAGG ACCACGACGGCGATTATAAGGATCACGACATCGACTACAAAGACGACGATGACAAGCATTAAAATGGACTCCAAAAACATTTTTTTAAATGTTCTG

KMT2E-CtermSanger-FwrP1: GGGCCACATTGTCCATTACC

KMT2E-CtermSanger-RevP1: CAACGAAAGAGCACTGGTGT

### Soluble Nuclear Extract Generation

Soluble nuclear extracts from HeLa-S3 cells (WT *vs. KMT2e*-3X FLAG) were generated, as previously described^85^. Briefly, cells were washed in PBS and resuspended in a low-salt buffer (LSB) containing 20 mM HEPES pH 7.9, 25% glycerol, 1.5 mM MgCl2, 2 mM EDTA, 1 mM DTT and Halt Protease and Phosphatase Inhibitor Cocktail. The cells were incubated on ice for 15 min to allow swelling, and non-ionic detergent NP-40 was added (final concentration of 0.75%), and the mixture was gently passed through a 21-gauge needle ten times to ensure lysis. The nuclei were collected by centrifugation at 1,100g for 5 min at 4 °C, and the supernatant discarded. The nuclei were washed 2X with LSB and resuspended in 500 µl of LSB. The pelleted nuclear volume (PNV) was calculated by subtracting 500 µl LSB from the total volume. Nuclei were then repelleted and resuspended in half PNV of LSB. An equivalent volume of high-salt buffer (20 mM HEPES pH 7.9, 25% glycerol, 1.5 mM MgCl2, 1.6 M NaCl, 1 mM DTT, Halt Protease and Phosphatase Inhibitor Cocktail) was added dropwise while vortexing at low speed to reach a final NaCl concentration of 400 mM. The samples were vortexed hard for 15 seconds, and then incubated at 4 °C with rotation for 1 h before being centrifuged at 21,000g for 10 min at 4 °C. The supernatant was collected as the soluble nuclear extract. For immunoprecipitations, the protein concentration was estimated by using a nanodrop. A small amount of input was saved for western blots, while the rest used for immunoprecipitations. Any remaining soluble nuclear extract was saved at -80°C, with only 1 freeze-thaw cycle allowed before sample was discarded.

### Immunoprecipitation

3 milligrams of WT or *KMT2E*-3xFLAG HeLa-S3 soluble nuclear extracts were utilized for all immunoprecipitations. 50 μL of Dynabeads Protein A or G (Rabbit or Mouse isotype, ThermoFisher 10002D/ 10004D) slurry was washed 2X in TBS-T and incubated with 5 ug of primary antibody for 1 hour at 4°C. Following 2 more washes, nuclear extract was added to the bead-antibody mixture, and low salt buffer used to dilute the incubation buffer to 150 mM NaCl before overnight rotation at 4 °C. The following morning, beads were washed 6X in high salt buffer (20 mM HEPES pH 7.9, 25% glycerol, 1.5 mM MgCl2, 500 mM NaCl, 1 mM DTT). For mass spectrometry analysis, beads were washed 2X in 1X TBS, pelleted, and frozen until mass spectrometry analysis performed. For western blotting, protein+antibody was eluted using 1X lamelli reducing load dye and boiled for 10 minutes prior to western blotting.

### Mass Spectrometry

For liquid chromatography tandem mass spectrometry analysis (LC-MS), proteins bound by antibody-bead conjugation were partially trypsinized (10ng/uL, Promega) for 3 hours. Supernatant was separated and reduced (10mM Dithiothreitol) and alkylated (30mM iodoacetamide). The supernatant was re-trypsinized overnight. Digestion was quenched by addition of 10% trifluoroacetic acid (TFA; Thermo Scientific 85183) and peptides were solid phase extracted^86^ prior to being analyzed by loading onto a reversed-phase nano-LC-MS/MS using a Fusion Ascend mass spectrometer operated in high/high acquisition mode. Peptides were separated using an EASY-Spray HPLC Column 25cm/75μm ID/2μm C18 column (ThermoFisher Scientific; CAT# ES902). Peptides were eluted using a gradient delivered at 350nL/min increasing from 1% Buffer B (0.1% formic acid in 80% acetonitrile) / 99% Buffer A (0.1% formic acid) to 30% Buffer B / 70% Buffer A, over 70 minutes (Neo Vanquish, Thermo Scientific). All solvents were LCMS grade (Optima, Fisher Scientific).

### Mass Spectrometry Data Processing and Analysis

Mass spectrometry data were searched against UniProts human protein database concatenated with common contaminant sequences^87^ using Thermo Proteome Discoverer/Sequest HT (v. 3.0) using full tryptic constraints. Oxidation of methionine residues and protein N-terminal acetylation were specified as variable modifications, and carbamidomethylation of cysteine residues was set as a fixed modification. False Discovery Rate (FDR) were set at 1% for both peptides and proteins. The experiment comprised five different conditions (WT-IgG, WT-FLAG, KMT2E-FLAG 150/300/500 mM wash), each run as three independent biological replicates. Quantitative data were processed using Perseus v. 1.6.15.0. Potential contaminants were removed prior to analysis. Raw abundance values were Log₂-transformed. Proteins were required to be quantified in at least three replicates in at least one condition to be included in downstream analyses. Missing values were imputed using a normal distribution (width = 0.3; downshift = 1.8). Statistical comparisons between only the WT-FLAG and KMT2E-FLAG 500 mM conditions were performed using a permutation-based two-sided t-test with an FDR threshold of 0.05.

### Cerebellum Granule Neuron Culturing

Culturing of primary cerebellar granule neurons was performed, as previously described^13^. Briefly, P7 pups were euthanized and cerebella were collected in freshly prepared HHGN media (1X HBSS, 2.5 mM HEPES pH 7.4, 35 mM glucose, and 4 mM sodium bicarbonate) on ice. Cerebella were washed 3X in ice cold HHGN media, then treated with trypsin for 15 minutes at 37°C. They were then washed 3X again in ice cold HHGN. A mixture of 200 μL of DNase in 5 mL of Basal Medium Eagle (BME, Thermo Fisher 21010046) was prepared, and 600 μL of this used to resuspend the washed cerebella by pipetting up and down > 20X with a P1000. 2 more mL of the BME/DNase was added, and sample incubated on ice 5 minutes. Then, supernatant was moved to a new tube, and remaining tissue resuspended again in another 1 mL of BME/DNase. Following 5 minute incubation on ice, the supernatant was combined with the previous supernatant. The cells in the supernatant were pelleted by spinning 500 x g for 5 minutes at room temperature. Cells were suspended in pre-warmed culture media (BME with 10% CGH Hyclone serum (FIsher SH103IH2540), 1X PenStrep, 1X GlutaMax, and 25 mM KCl; filtered) and counted using a hemocytometer. 15-20 million cells were plated per 10 cm dish (pre-coated with 1X Poly-L-Ornithine). Half media changes were performed every 3 days, and neurons considered mature at DIV7. For *Kmt2e* mutant mice, individual cerebella remained separate through isolation and culturing, with a small amount of tissue retained for genotyping using the protocol described above.

### MB-3 Treatment of Neurons

Butyrolactone 3 (MB-3, MedChemExpress HY-129039) was reconstituted in a 5% DMSO:95% of a 20% HP-β-CD (MedChemExpress, HY-101103) mixture, at a stock concentration of 2 mg/ml (10.85 mM). A dilution series was performed to identify the optimal concentration to treat cultured neurons for 24 hours, using sodium butyrate as a negative control. Ultimately, 50 uM was utilized for treating cells for 24 hours. Neurons were monitored visually for cytotoxicity, and vehicle control was used (5% DMSO:95% of a 20% HP-β-CD).

### Antibodies

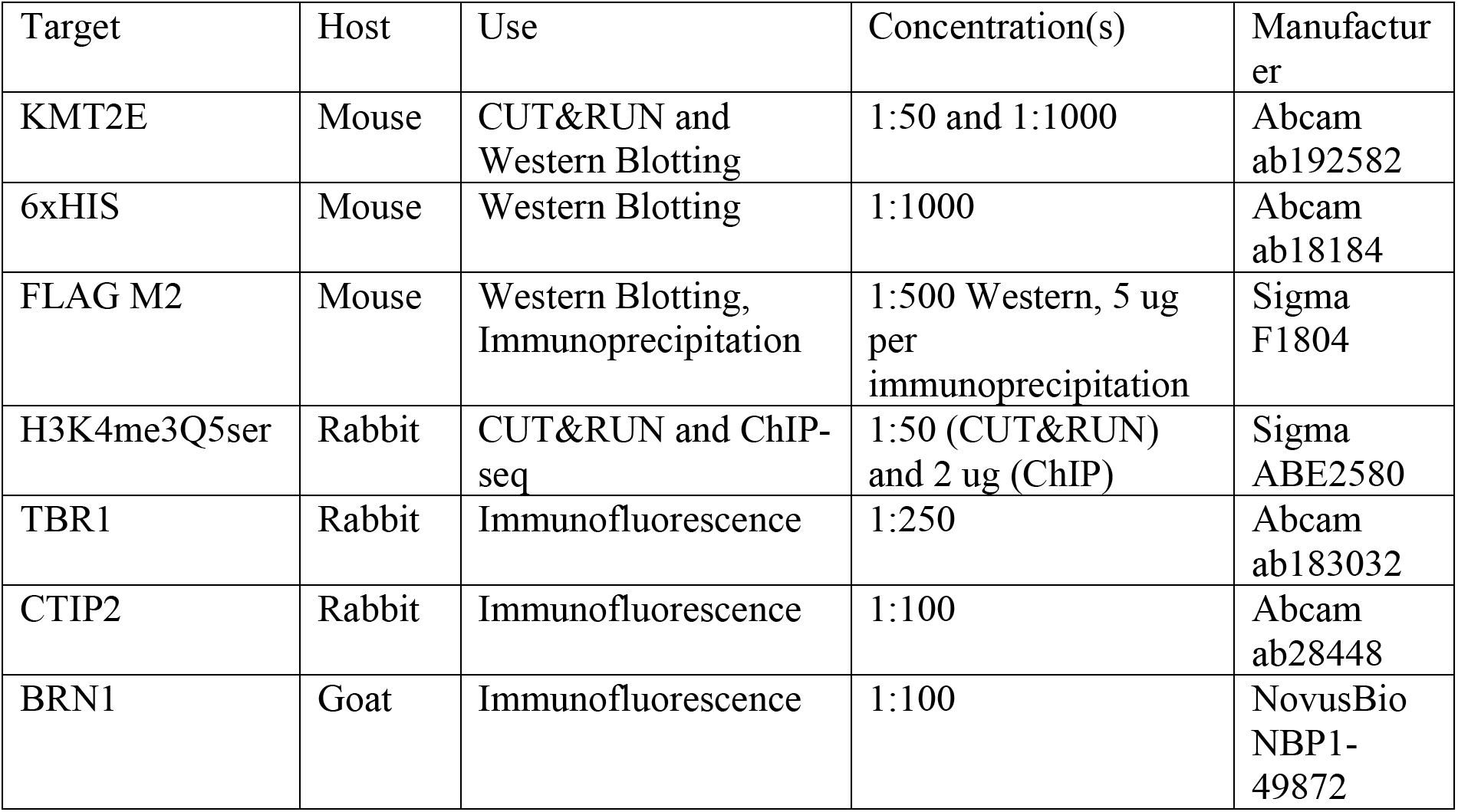
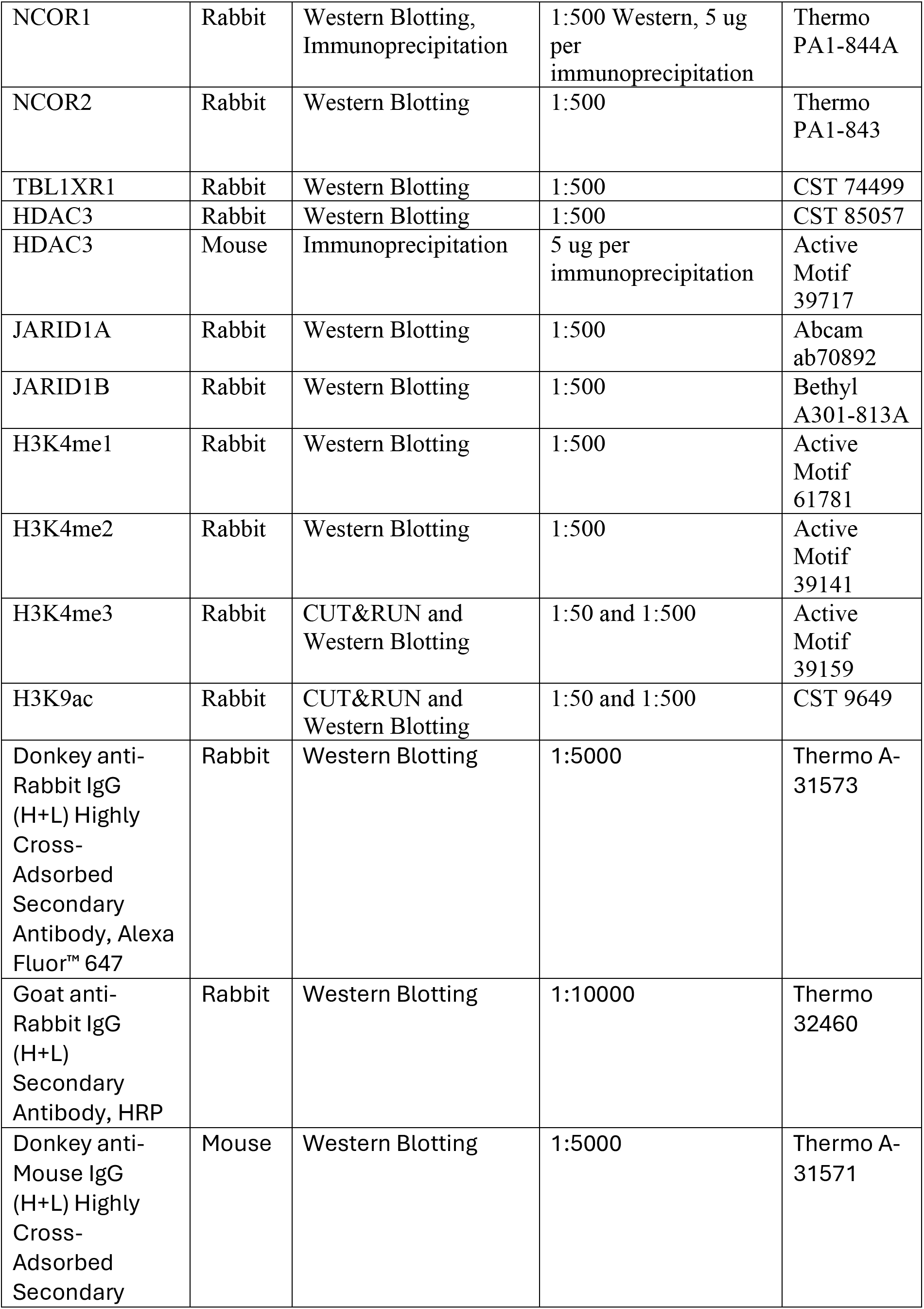
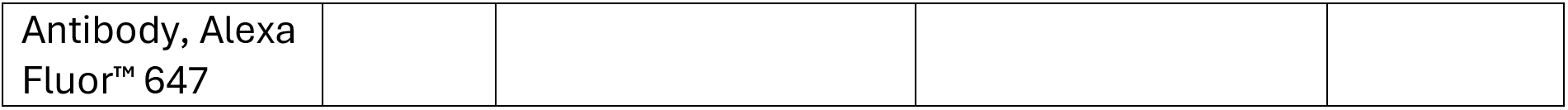

## ACKNOWLEDGEMENTS

We would like to thank members of the Maze lab for helpful discussions on this study. This work was partially supported by grants from the National Institutes of Health: R01 MH116900 (I.M.), F32 MH140478 (B.H.W.), F32 MH126534 and K01 MH139990 (J.C.O’C.), R21 NS130319 (S.G.M.), F31 NS132558 (A.M.C.), F99 NS139541 (A.M.C.), F30 AG090096 (W.C.), 3T32ADA007135 (N.A.), as well as funds from the Brain and Behavior Research Foundation (J.C.O’C.), and the Howard Hughes Medical Institute (I.M.). Additional support was provided by a Cindy Silvian Foundation Grant to S.G.M, and a Simons Collaboration on Plasticity and the Aging Brain Fellows-to-Faculty Award (Z.C.W). All schematics were created with Biorender.com. Crystallography data was collected with aid from Dr. Shilong Fan, X-ray facility manager at the Advanced Innovation Center for Structural Biology, Tsinghua University. The crystallized peptide, H3(1–7)K4me3Q5ser, was generously provided by Dr. Xiang Li’s laboratory at the University of Hong Kong. Experiments were supported by the Stem Cell Engineering Core (RRID:SCR_027503) at the Icahn School of Medicine at Mount Sinai. Electron Microscopy tissue preparation and ultrastructural imaging were performed at The Microscopy and Advanced Bioimaging Core at the Icahn School of Medicine at Mount Sinai.

## AUTHOR CONTRIBUTIONS

J.C.O’C., B.H.W., and I.M. conceived of the study, designed the experiments and interpreted the data. J.C.O’C., W.C., R.I., N.A., and E.A. performed behavioral experiments. J.C.O’C., Z.C.W and T.S. performed and analyzed kainic acid seizure susceptibility experiments. J.C.O’C. and B.H.W. performed all molecular experiments. J.C.O’C., B.H.W., A.R., and L.S. performed all bioinformatic analyses. E.B. and S.G.M. created the mouse line and performed validation. B.H.W. performed CRISPR knock-in and validation, as well as all immunoprecipitations and co-immunoprecipitations. C.P. and H.M. performed LC-MS/MS runs and analyses. J.C.O’C. performed all primary neuronal culturing. C.Y. and H.L. performed x-ray crystallography, isothermal titration calorimetry, and all related annotation and analyses. B.L. and T.W.M. provided histone H3 peptides for all assays. C.A.S. and M.T.B. performed peptide array experiment and analysis. V.P. and B.D.B performed electrophysiology and analyses. J.C.O’C., B.H.W., M.C., and B.C. performed cloning. J.C.O’C., S.D. and A.M.C performed immunofluorescence and electron microscopy and related analyses. J.C.O’C., B.H.W., and I.M. wrote and edited the manuscript, with input from all co-authors where appropriate.

## COMPETING INTERESTS

The authors declare no competing interests.

## DATA AVAILABILITY

The genomics data generated in this study have been deposited in the National Center for Biotechnology Information Gene Expression Omnibus (GEO) database under the SuperSeries GSE339280. All mass spectrometry-based proteomics data have been deposited to the ProteomeXchange Consortium via the PRIDE partner repository with the datasets identified as PXD081270. The atomic coordinates and structure factors have been deposited in the Protein Data Bank (PDB) under PDB ID 45YW. We declare that the data supporting findings for this study are available within the article and Supplementary Information. Related data including raw microscopy images are available from the corresponding author upon reasonable request. No restrictions on data availability apply.

## CODE AVAILABILITY

Related code is available from the corresponding author on reasonable request.

## EXTENDED DATA TABLES/FIGURES AND LEGENDS

**Extended Data Table 1.**
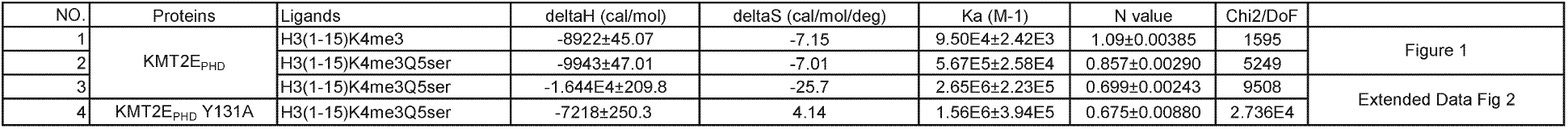
ITC statistics.

**Extended Data Table 2.** X-ray crystallography data collection and refinement statistics.

| Crystal | KMT2E<sub>PHD</sub>-H3K4me3Q5ser |
| --- | --- |
| **Data collection** |  |
| Wavelength(Å) | 0.9792 |
| Space group | P1 |
| Cell dimensions |  |
| a, b, c (Å) | 33.9, 38.0, 43.3 |
| α, β, γ (°) | 65.2, 84.9, 71.0 |
| Resolution (Å) | 50 - 2.20 (2.24 - 2.20) |
| No. of reflections | 9105 (427) |
| R<sub>merge</sub> (%) | 9.3 (23.9) |
| R<sub>pim</sub> (%) | 5.9 (15.8) |
| I/σ (I) | 18.8 (4.9) |
| CC1/2 (%) | 97.0 (94.2) |
| Completeness (%) | 97.9 (93.4) |
| Redundancy | 3.5 (3.0) |
| **Refinement (F>0)** |  |
| R<sub>work</sub>/R<sub>free</sub> (%) | 15.5/ 20.4 |
| No. of non-H atoms |  |
| Protein | 1421 |
| Peptide | 46 |
| Water | 101 |
| Average B-factor (Å<sup>2</sup>) |  |
| Protein | 36.01 |
| Peptide | 44.64 |
| Water | 38.91 |
| RMSD bonds (Å) | 0.006 |
| RMSD angels (°) | 0.844 |
| **Ramachandran plot** |  |
| Favored (%) | 98.80 |
| Allowed (%) | 1.20 |
| Outliers (%) | 0.00 |

**Extended Data Figure 1.**
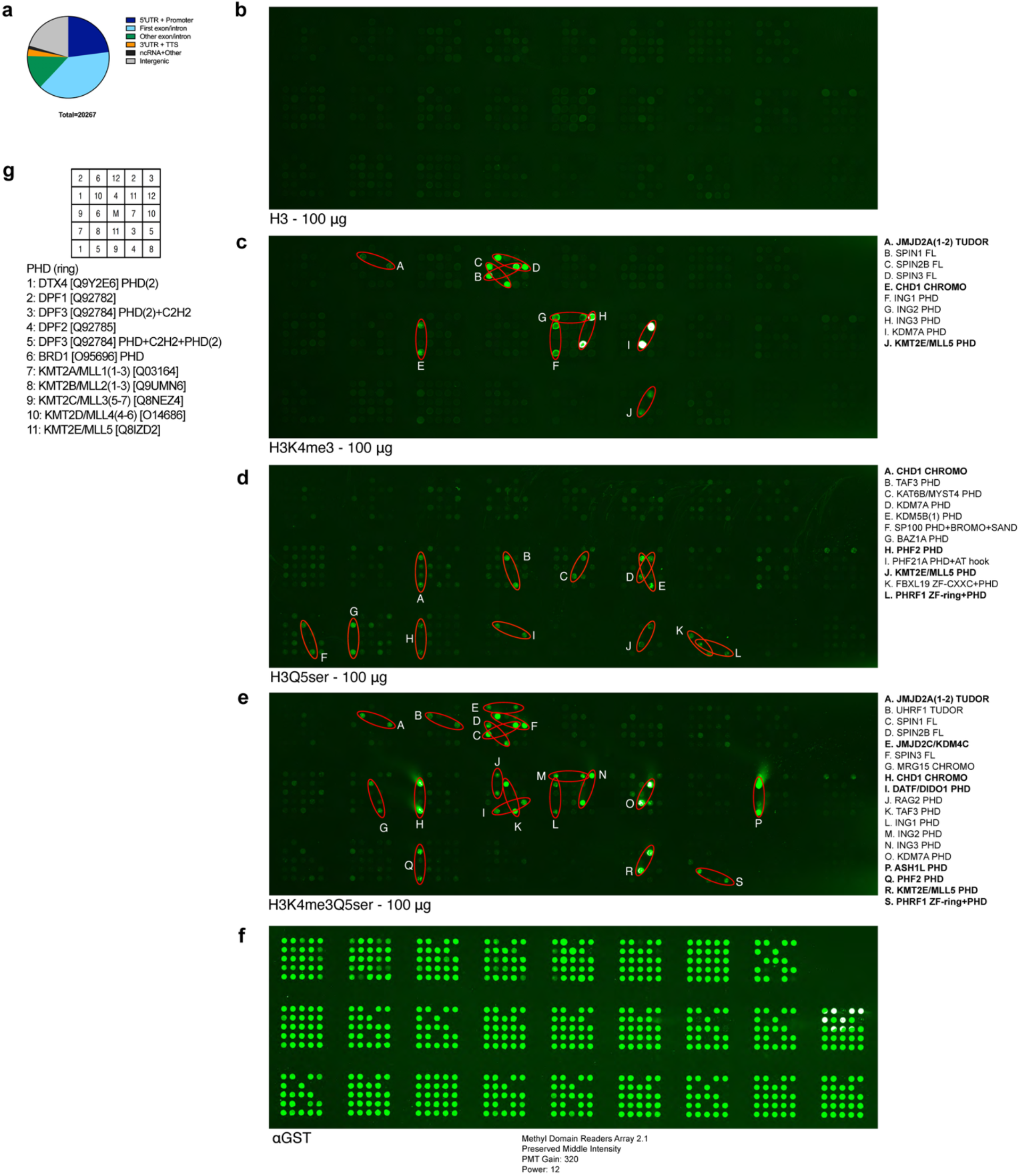
High-throughput detection of H3K4me3Q5ser reader interactions using protein domain microarrays. **(a**) Distribution of H3K4me3Q5ser genomic enrichment in E12.5 mouse forebrain tissues. (**b-f**) Methyl domain reader arrays probed with (**b**) H3_1-10_, (**c**) H3K4me3_1-10_, (**d**) H3Q5ser_1-10_, or (**e**) H3K4me3Q5ser_1-10_ peptides *vs.* (**f**) an anti-GST loading control. The panel in (**f**) shows the position of all arrayed recombinant proteins. Red circles indicate domain *readers* run in duplicate with significant binding, as indicated by the corresponding text on the right. Bolded text indicates proteins previously implicated in NDDs. Note, the full list of proteins contained within the arrays is provided in Vaughan et al, 2020^78^. (**g**) All PHD fingers probed and displayed in Fig. 1e.

**Extended Data Figure 2.**
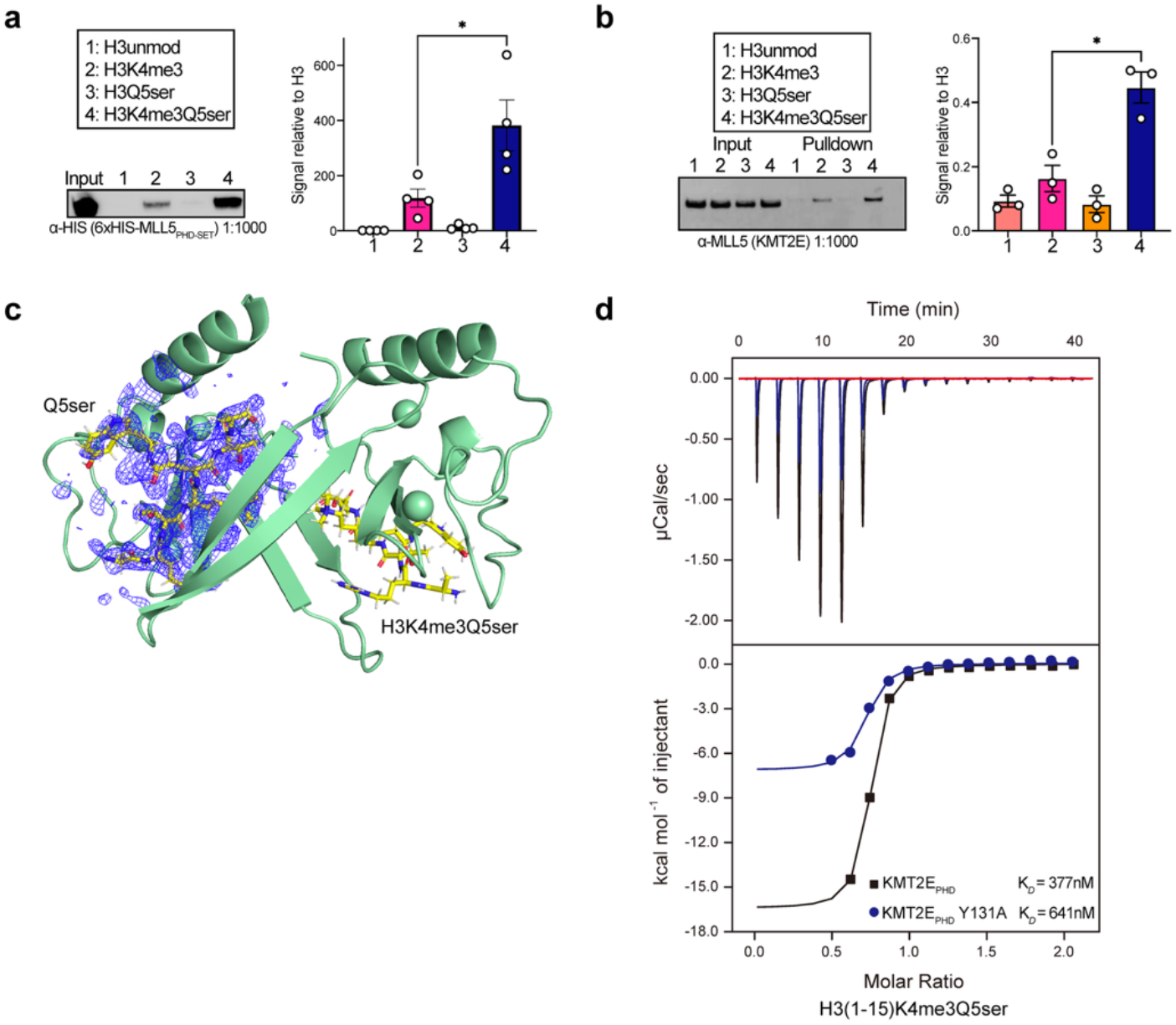
Validation of KMT2E-H3K4me3Q5ser interactions. (**a**) Peptide pulldown validations using recombinant tagged (6xHIS) KMT2E_PHD-SET_, followed by immunoblotting for the His tag. *n* = 4/group. One-way ANOVA: *p* = 0.0004; Tukey’s *post-hoc*, H3K4me3 vs. H3K4me3Q5ser: \**p* = 0.0117. (**b**) H3_1-10_ peptide immunoprecipitations from HeLa nuclear extracts, followed by immunoblotting for KMT2E. *n* = 3/group. One-way ANOVA: *p* = 0.0003; Tukey’s *post-hoc* H3K4me3 vs. H3K4me3Q5ser: \*\**p* = 0.022. (**c**) Electron density of the H3K4me3Q5ser peptide in blue mesh. The green ribbon is the KMT2E_PHD_ dimer. (**d**) Titration and fitting curve for H3K4me3Q5ser peptides titrated into KMT2E_PHD_ WT *vs.* KMT2E_PHD_ Y131A mutant, showing ∼1.7x reduction in *K_D_.* Uncropped blots are provided in **Supplementary Figure 1**. Bar plots are mean ± SEM.

**Extended Data Figure 3.**
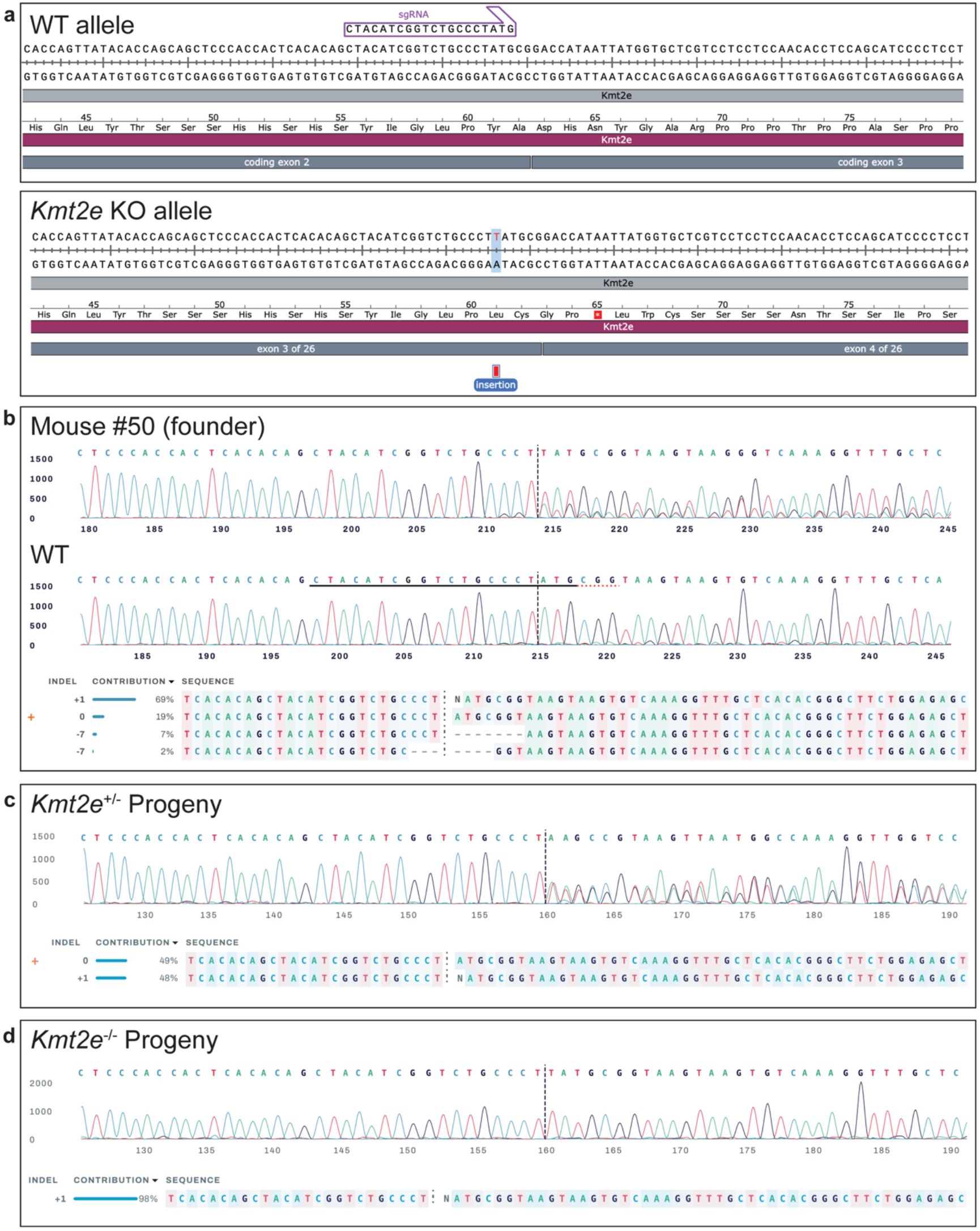
Generation of *Kmt2e* KO mutant mice. (**a**) Schematic alignment of the wild-type (WT) genomic sequence and the edited Kmt2e locus in Mouse #50 (founder). The sgRNA targeting sequence is highlighted in purple. In the edited KO allele, a 1-bp thymidine (T) insertion shifts the reading frame starting at codon 61. This frameshift changes the downstream amino acid sequence and introduces a premature stop codon (red asterisk, TAA) four codons downstream of the insertion. Note: exon 3 of the murine *Kmt2e* locus is coding exon 2. (**b**) Sanger sequencing chromatogram from Mouse #50 (founder, top) compared to a control sample (middle). Below, the Synthego Inference of CRISPR Edits (ICE) deconvolution analysis illustrates the indel distribution, highlighting the target cleavage site (vertical dashed line) and the presence of the +1 frameshift allele (69% contribution) alongside the unedited allele (19% contribution). (**c**) Representative Sanger sequencing chromatogram from a heterozygous (+/-) progeny. ICE deconvolution analysis illustrates the presence of the +1 frameshift allele (48% contribution) alongside the unedited allele (49% contribution). (**d**) Representative Sanger sequencing chromatogram and ICE deconvolution from a homozygous Kmt2e KO (-/-) progeny, showing 98% contribution of the edited locus.

**Extended Data Figure 4.**
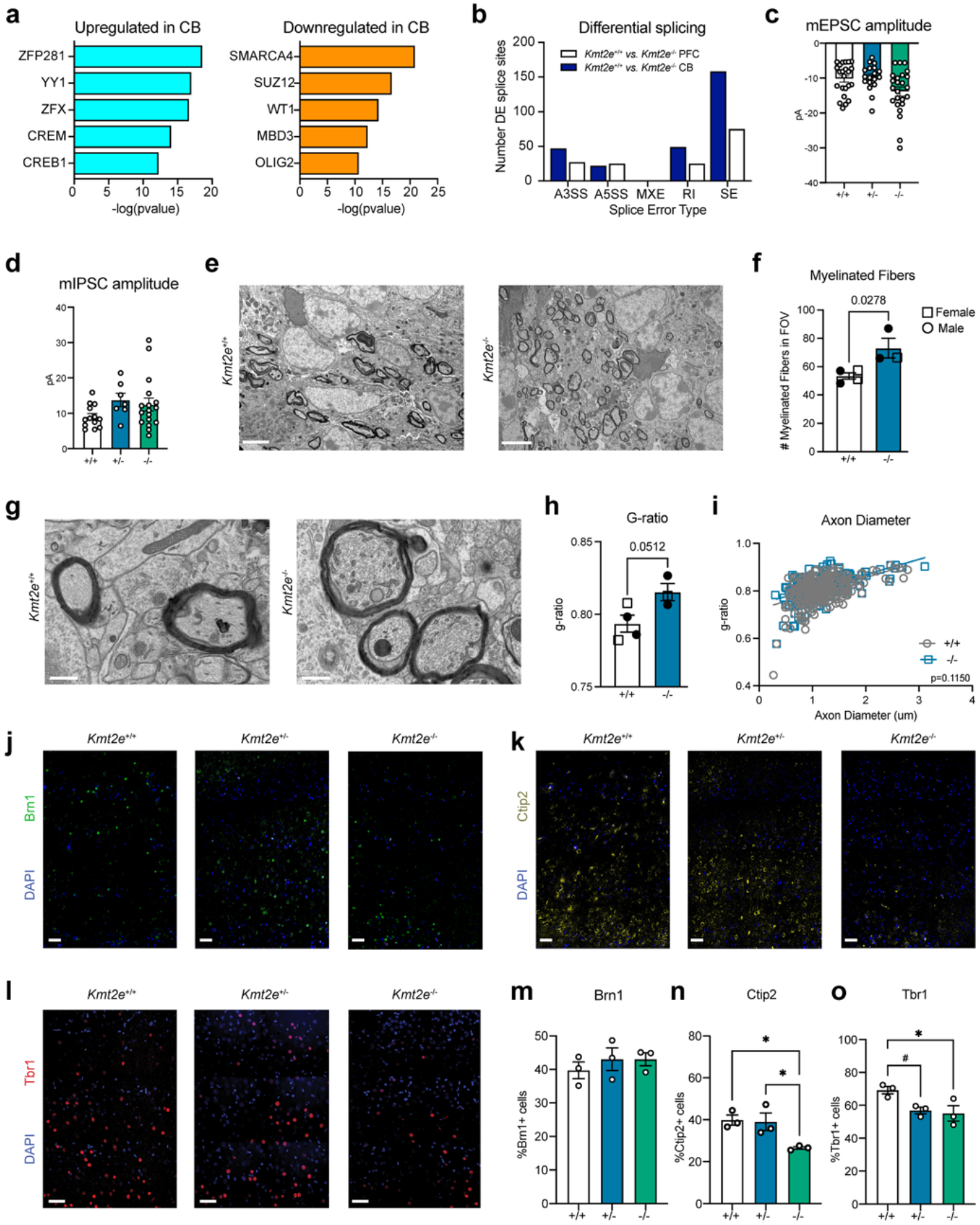
Cellular, physiological, and molecular outcomes in Kmt2e KO brain tissues. (**a**) ChEA ontology analysis for upregulated and downregulated DEGs in WT *vs.* KO cerebellum. (*p* < 0.05). (**b**) Number of differential alternative splicing events for junction counts and exon counts using rMATS analysis. A3SS, alternative 3’ splice site; A5SS, alternative 5’ splice site; MXE, mutually exclusive exons; RI, retained intron; SE, skipped exon. (**c**) Cell mean of mEPSC amplitude for WT (*n* = 24), HET (*n* = 19), and KO (*n* = 24) cerebellar granule neurons. Kruskal-Wallis test: *p* = 0.0344; Dunn’s *post-hoc*: WT *vs.* KO, *p* = 0.0628. (**d**) Cell mean of mIPSC amplitude for WT (*n* = 13), HET (*n* = 7), and KO (*n* = 17) cerebellar granule neurons. Kruskal-Wallis test: *p* = 0.1760. (**e**) Representative transmission electron microscopy images of CB projections. Scale bars, 4 μm. (**f**) Quantification of myelinated fibers in the field of view (FOV). Unpaired two-tailed Student’s t-test: *p* = 0.0278. (**g**) Representative electron micrographs showing myelinated axons in CB. Scale bars, 600 nm. (**h**) Increased *g*-ratio of axons in KO CB, indicating a decrease in myelin sheath thickness. Unpaired two-tailed Student’s t-test: *p* = 0.0512. (**i**) Scatter plot of *g*-ratio as a function of axon diameter. WT (grey circles), KO (blue squares). For (f) and (h), WT, *n* = 4; KO, *n* = 3. (**j-l**) Representative images of staining of cortical layer markers (**j**) Brn1, (**k**) Ctip2, and (**l**) Tbr1 in P14 male mPFC. (**m-o**) Quantification of (**m**) Brn1; one-way ANOVA: *p* = 0.5101; (**n**) Ctip2; one-way ANOVA: *p* = 0.0264; (**o**) Tbr1; one-way ANOVA, *p* = 0.0409. Tukey’s *post hoc*: \**p* < 0.05, #*p* = 0.06. For m-o, *n* = 3/genotype. Scale bars, 50 um. Bar plots are presented as mean ± SEM.

**Extended Data Figure 5.**
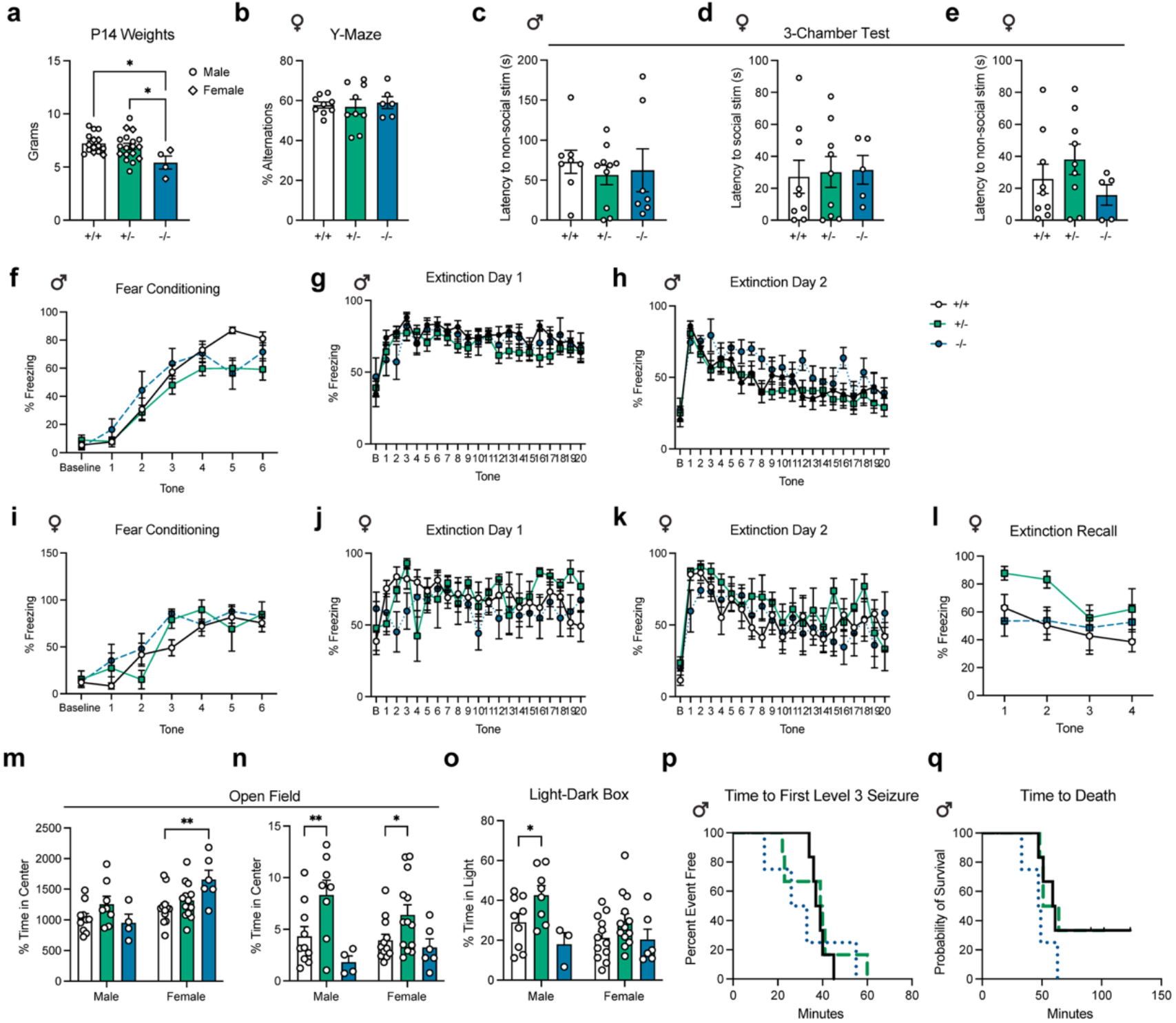
Kmt2e mutant mice mimic sex-dependent endophenotypes of ODLURO syndrome. (**a**) Weights for P14 mice, averaged per litter for males (WT, *n* = 9; HET, *n* = 11; KO, *n* = 2) and females (WT, *n* = 6; HET, *n* = 7; KO, *n* = 2). One-way ANOVA: *p* = 0.240; Tukey’s *post-hoc*, \**p* < 0.05. (**b**) Female (WT, *n* = 9; HET, *n* = 9; KO, *n* = 6) mice on the Y-maze. One-way ANOVA: *p* = 0.8896. (**c**) No differences for male (WT, *n* = 8; HET, *n* = 10; KO, *n* = 7) mice in approach of a novel non-social stimulus on the 3-Chamber Social Test. One-way ANOVA: *p* = 0.7863. (**d-e**) No differences for female (WT, *n* = 9; HET, *n* = 9; KO, *n* = 5) mice in approach of a novel (**d**) social or (**e**) non-social stimulus on the 3-Chamber Social Test. One-way ANOVA, (d) *p* = 0.9567; (e) *p* = 0.3012. (**f**) Acquisition of auditory fear conditioning in male mice. Two-way RM ANOVA: time: *p* < 0.0001; genotype, *p* = 0.0733; interaction: *p* = 0.0483. Tukey’s *post-hoc*: tone_5_: WT *vs.* HET, *p* = 0.0004; tone_6_: WT *vs.* HET, *p* = 0.0549. (**g-h**) No effects of genotype for extinction of fear conditioning in the (**g**) training context nor in a (**h**) novel context. For (f-h), WT, *n* = 12; HET, *n* = 19; KO, *n* = 8. (**i-l**) No effects of genotype for females on (**i**) acquisition of auditory fear conditioning, extinction in the (**j**) training or (**k**) novel context, or in (**l**) extinction recall. For (i-l), WT, *n* = 7; HET, *n* = 4; KO, *n* = 5. (**m**) Total distance on the open field test. Two-way ANOVA: sex: *p* = 0.001; genotype, *p* = 0.1088; interaction: *p* = 0.0256. Tukey’s *post-hoc*: \*\**p* < 0.01. (**n**) Duration of time spent in the center on the open field test. Two-way ANOVA: sex: *p* = 0.7395; genotype, *p* = 0.0002; interaction: *p* = 0.3555. Tukey’s *post-hoc*: \**p* < 0.05, \*\**p* < 0.01. For (m-n), Males: WT, *n* = 10; HET, *n* = 8; KO, *n* = 4. Females: WT, *n* = 12; HET, *n* = 13; KO, *n* = 6. (**o**) Duration of time spent in the light on the light-dark box. Two-way ANOVA: sex: *p* = 0.1273; genotype, *p* = 0.0034; interaction: *p* = 0.3550. Tukey’s *post-hoc*: \**p* < 0.05. Males: WT, *n* = 9; HET, *n* = 8; KO, *n* = 3. Females: WT, *n* = 12; HET, *n* = 13; KO, *n* = 6. (**p-q**) Survival curves showing the difference between genotypes using the log-rank test for time to (**p**) first level 3 seizure (χ² = 0.6495, df = 2, *p* = 0.7227) and (**q**) death (χ² = 4.085, df = 2, *p* = 0.1297) in male mice. For p-q: WT (*n* = 6), HET (*n* = 6), KO (*n* = 4). Data are presented as mean ± SEM.

**Extended Data Figure 6.**
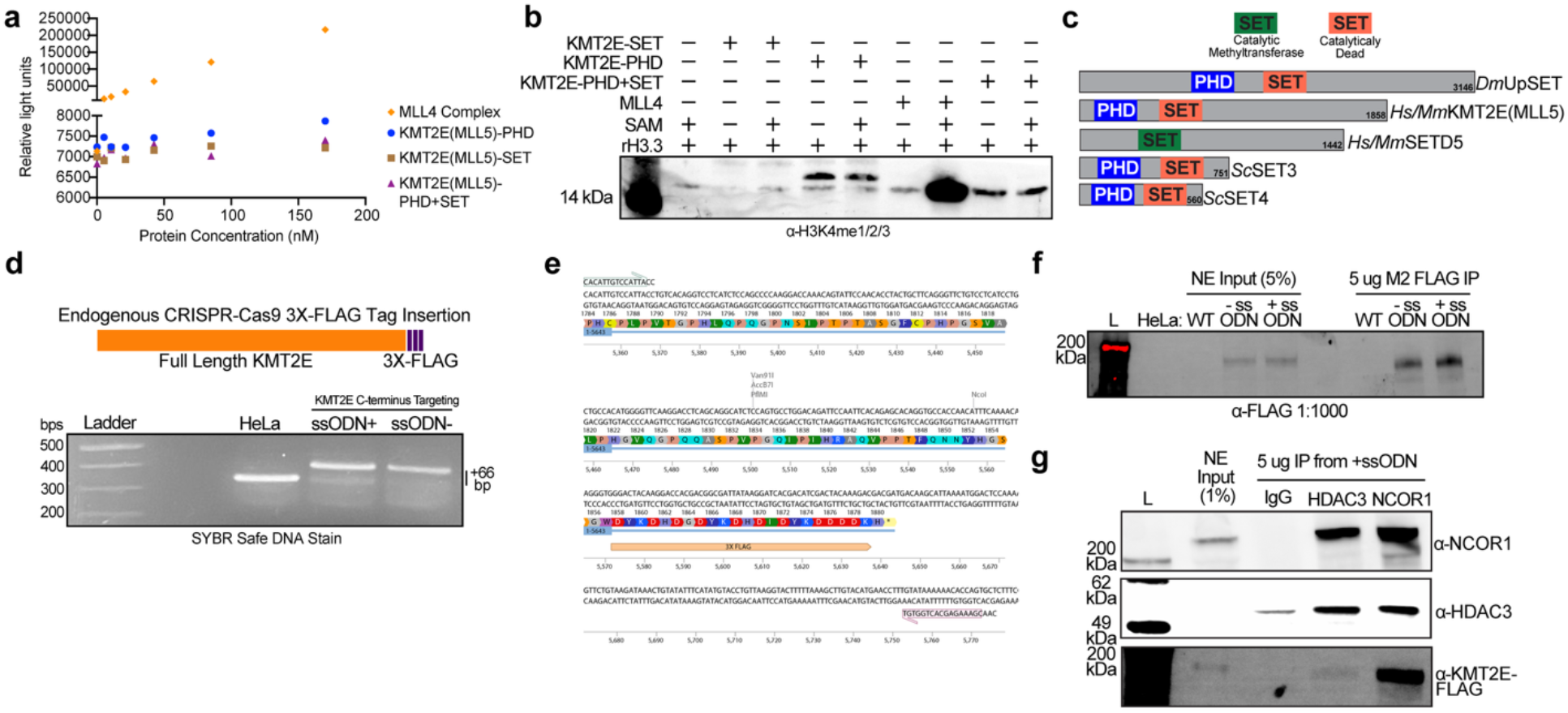
Catalytically inactive KMT2E interacts with the NCoR/HDAC3 repressor complex. (**a**) KMT2E PHD, SET, and PHD-SET domains displayed minimal activity in methyltransferase activity assays relative to the MLL4 complex, indicating lack of enzymatic activity. (**b**) *In vitro* methyltransferase reactions with recombinant H3.3 (rH3.3) comparing KMT2E-SET, KMT2E-PHD, KMT2E-PHD+SET *vs.* MLL4 complex, blotted for H3K4 mono-, di-, and tri-methylation. (**c**) Comparison of KMT2E and homologous proteins across species, with PHD and SET (active, green; inactive, orange) domains notated. *Dm*, *Drosophila melanogaster; Hs, Homo sapiens; Mm, Mus musculus; Sc, Saccharomyces cerevisiae.* (**d**) CRISPR/Cas9-mediated knock-in strategy of the endogenous KMT2E locus with a 3xFLAG tag at the C-terminus (top), which was confirmed by altered amplicon size in HeLa cells with single-stranded oligodeoxynucleotide (ssODN) targeted to the positive (+) or negative (-) strand. (**e**) Sequence validation of the KMT2E C-terminus showing in-frame incorporation of the 3xFLAG tag. (**f**) Immunoprecipitation of tagged KMT2E from nuclear extracts (NE) of WT and ssODN+ targeted HeLa cells, immunoblotted for the FLAG tag. (**g**) Reverse co-immunoprecipitation for the indicated proteins or IgG control from HeLa ssODN+ targeted HeLa cells. Uncropped blots are provided in **Supplementary Figure 1**.

**Extended Data Figure 7.**
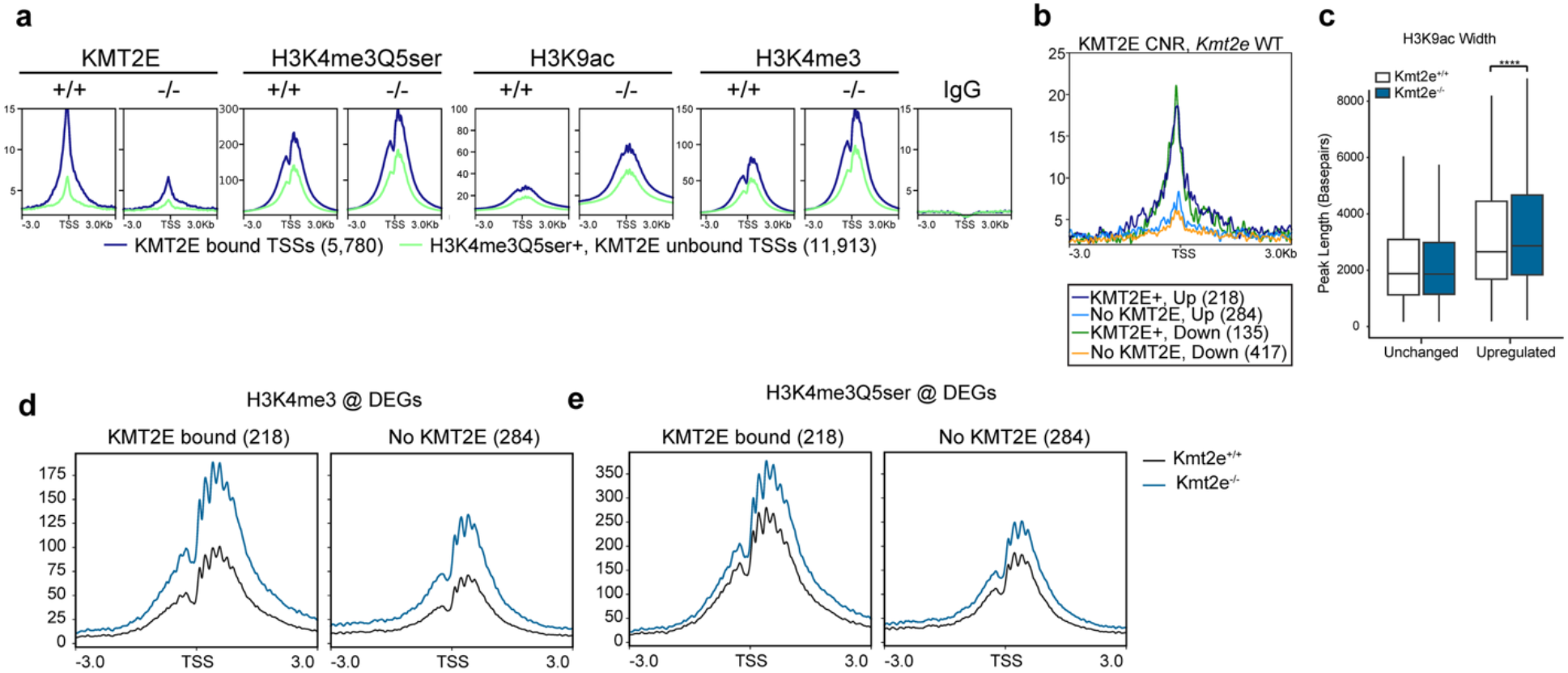
Loss of KMT2E alters permissive histone PTM levels at differentially expressed genes. (**a**) CUT&RUN-seq profiles for the indicated targets (KMT2E, H3K4me3Q5ser, H3K9ac, IgG) at KMT2E-bound (KMT2E+, blue line) *vs.* unbound (KMT2E-, green line) TSSs. (**b**) CUT&RUN-seq profiles for KMT2E at upregulated *vs.* downregulated DEGs in WT vs. KO CB. (**c**) H3K9ac peak width are increased in DEGs upregulated in KO CB. *n* = 4/genotype. Wilcoxon rank-sum test, \*\*\*\**p* < 2.2 x 10^-16^. Box plot shows median (center line), interquartile range (box), and minimum-to-maximum values (whiskers). (**d**) CUT&RUN-seq profiles for H3K4me3 at TSSs of upregulated DEGs, separated by KMT2E occupancy, showing H3K4me3 levels are higher in KO CB. (**e**) CUT&RUN-seq profiles for H3K4me3Q5ser at TSSs of DEGs, separated by KMT2E occupancy, showing H3K4me3Q5ser levels are higher in KO CB.

**Extended Data Figure 8.**
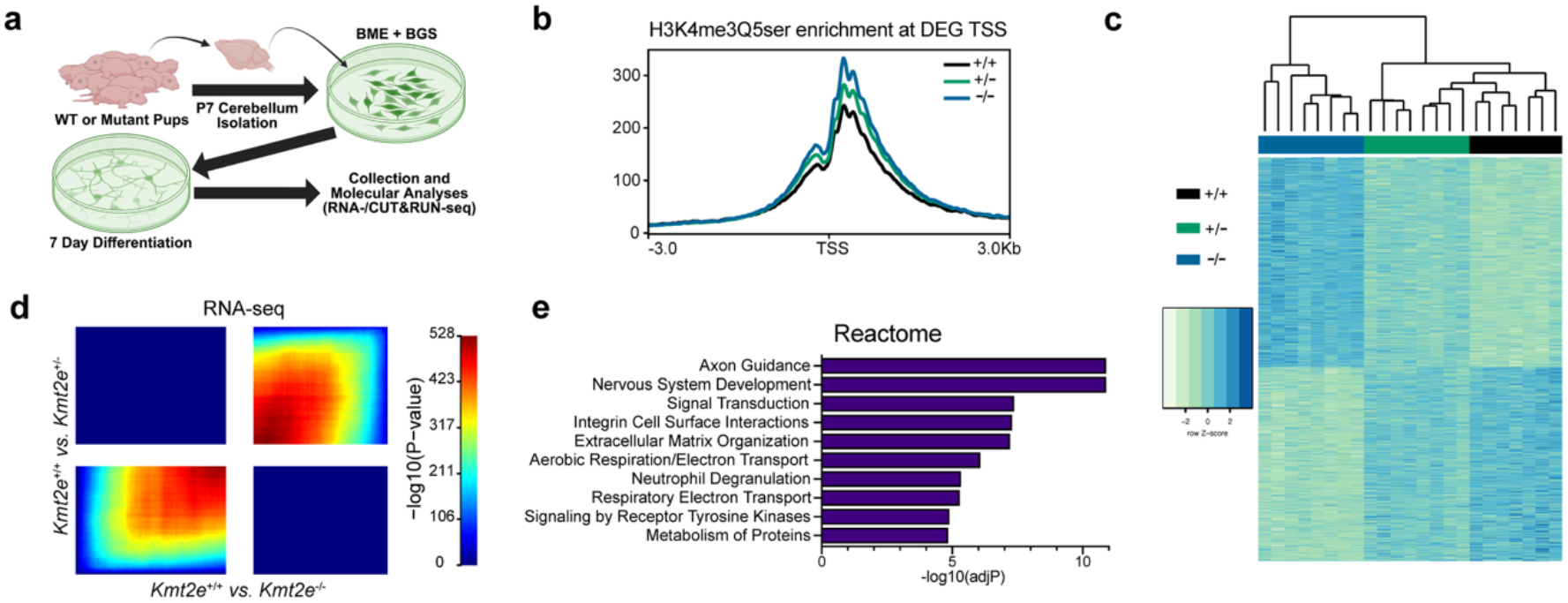
KMT2E haploinsufficiency dysregulates H3K4me3Q5ser and gene expression profiles. **(a**) Schematic showing primary CB granule neuron culture protocol from Kmt2e WT, HET and KO P7 pups. (**b**) Plot profiles from H3K4me3Q5ser CUT&RUN-seq at TSSs of WT *vs.* KO DEGs in CB. *n* = 4/genotype. (**c**) Heatmap of DEGs (DESeq2, *padj* < 0.05), with hierarchical clustering. *n* = 6/genotype. (**d**) Threshold-free comparison using RRHO, showing concordant gene expression profiles between WT vs. HET and WT vs. KO CB. (**e**) Reactome terms enriched for DEGs (*padj* < 0.05).

**Extended Data Figure 9.**
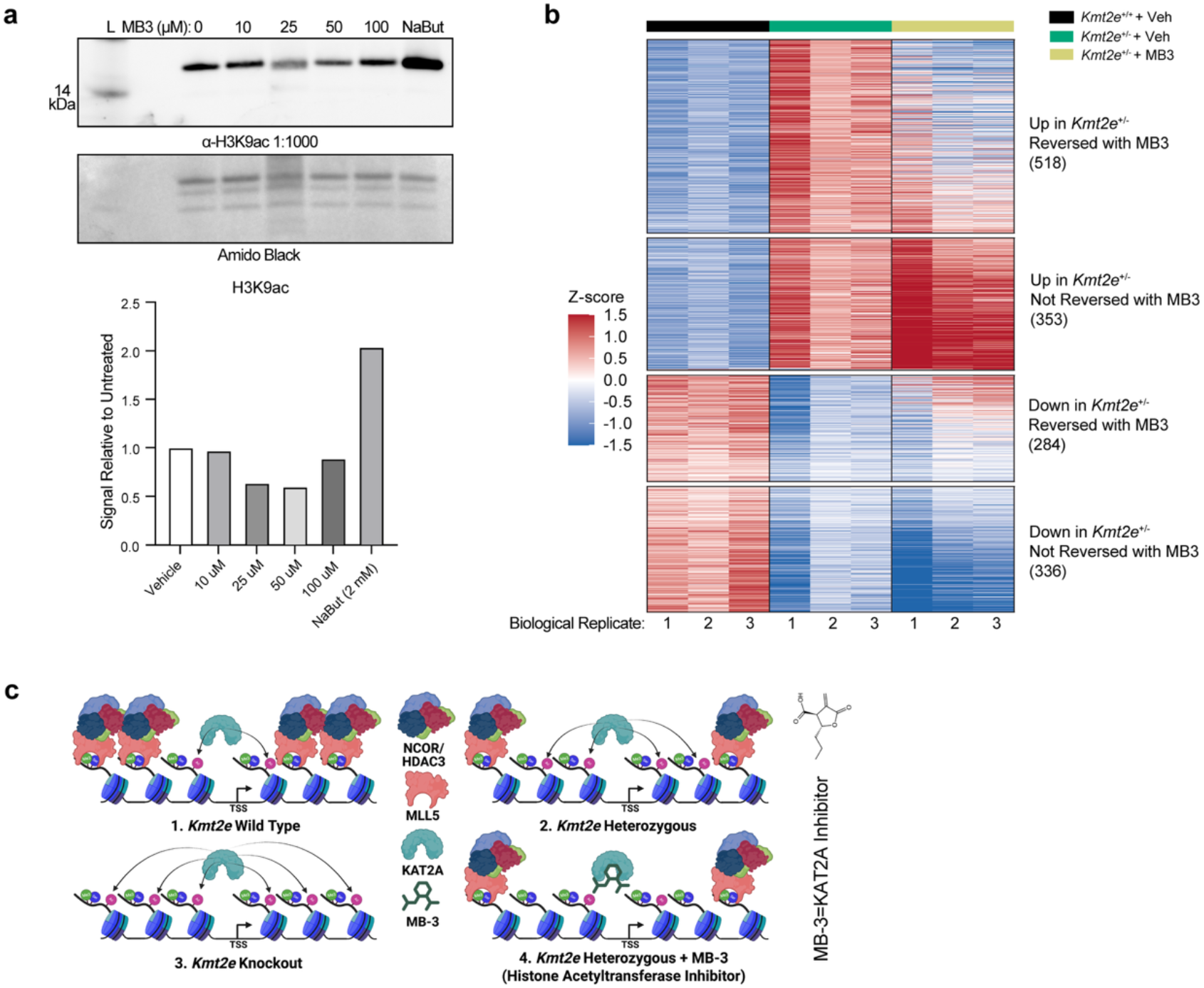
MB-3 treatment partially reverses transcriptional dysregulation in Kmt2e haploinsufficient neurons. (**a**) H3K9ac levels in primary neuronal cultures following MB-3 treatment. The histone deacetylase inhibitor sodium butyrate (NaBut) was used as a positive control. (**b**) Heatmap of DEGs across WT, HET, and HET+MB-3 conditions, showing clusters of genes upregulated or downregulated in HET that are either reversed or unchanged following MB-3 treatment. *n* = 3/group. (**c**) Schematic depicting the proposed mechanism by which KMT2E regulates chromatin dynamics. In WT cells, KMT2E is recruited to TSSs by H3K4me3Q5ser, where it recruits the NCoR/HDAC3 complex to restrict H3K9ac levels. Loss of KMT2E in HET and KO results in reduced NCoR/HDAC3 recruitment and increased H3K9ac levels, likely via KAT2A. Treatment with the KAT2A inhibitor MB-3 restores H3K9ac levels and gene expression in Kmt2e mutants. Uncropped blots are provided in **Supplementary Figure 1**.

## SUPPLEMENTARY MATERIALS

**Supplementary Figure 1:**
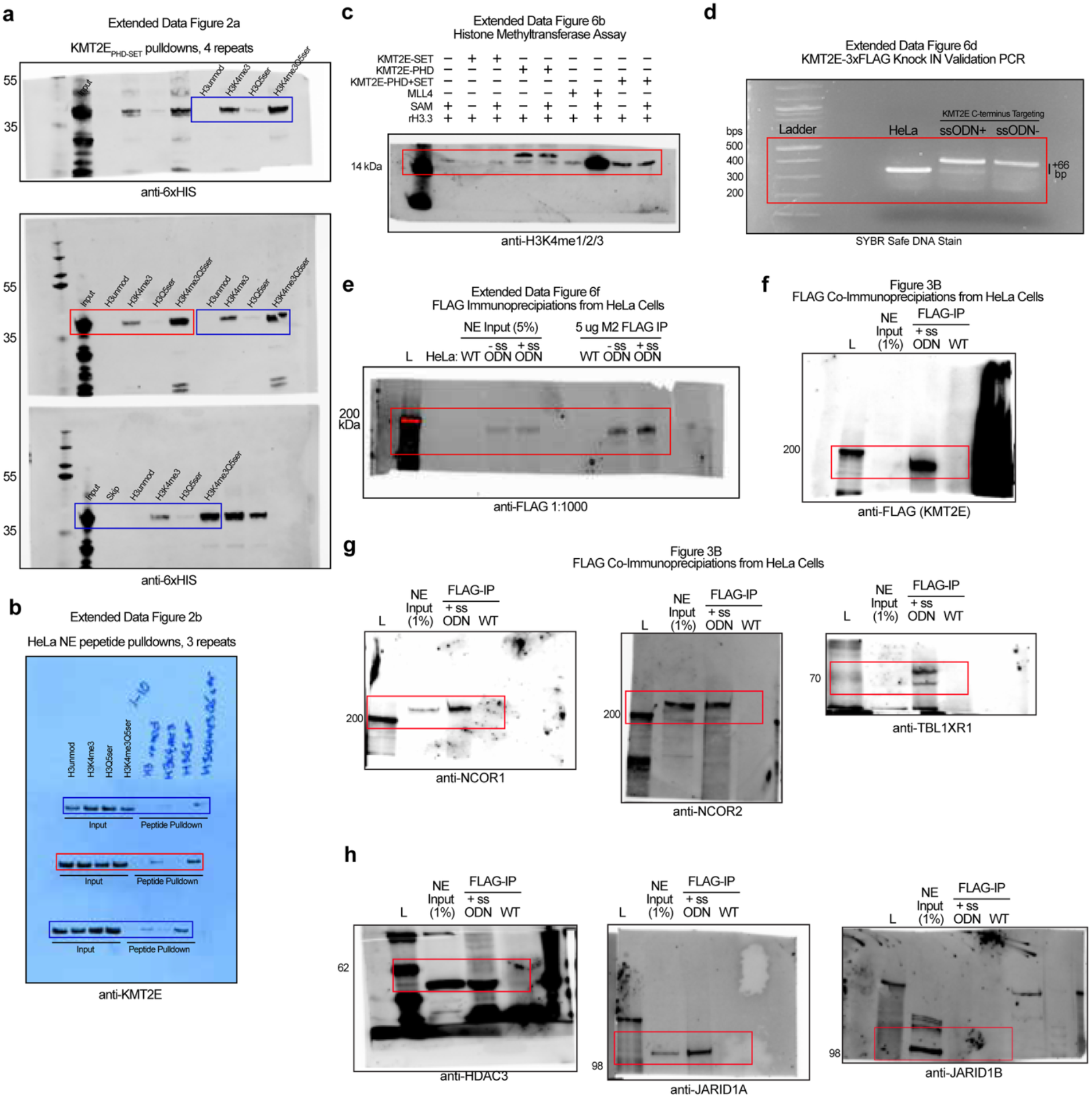
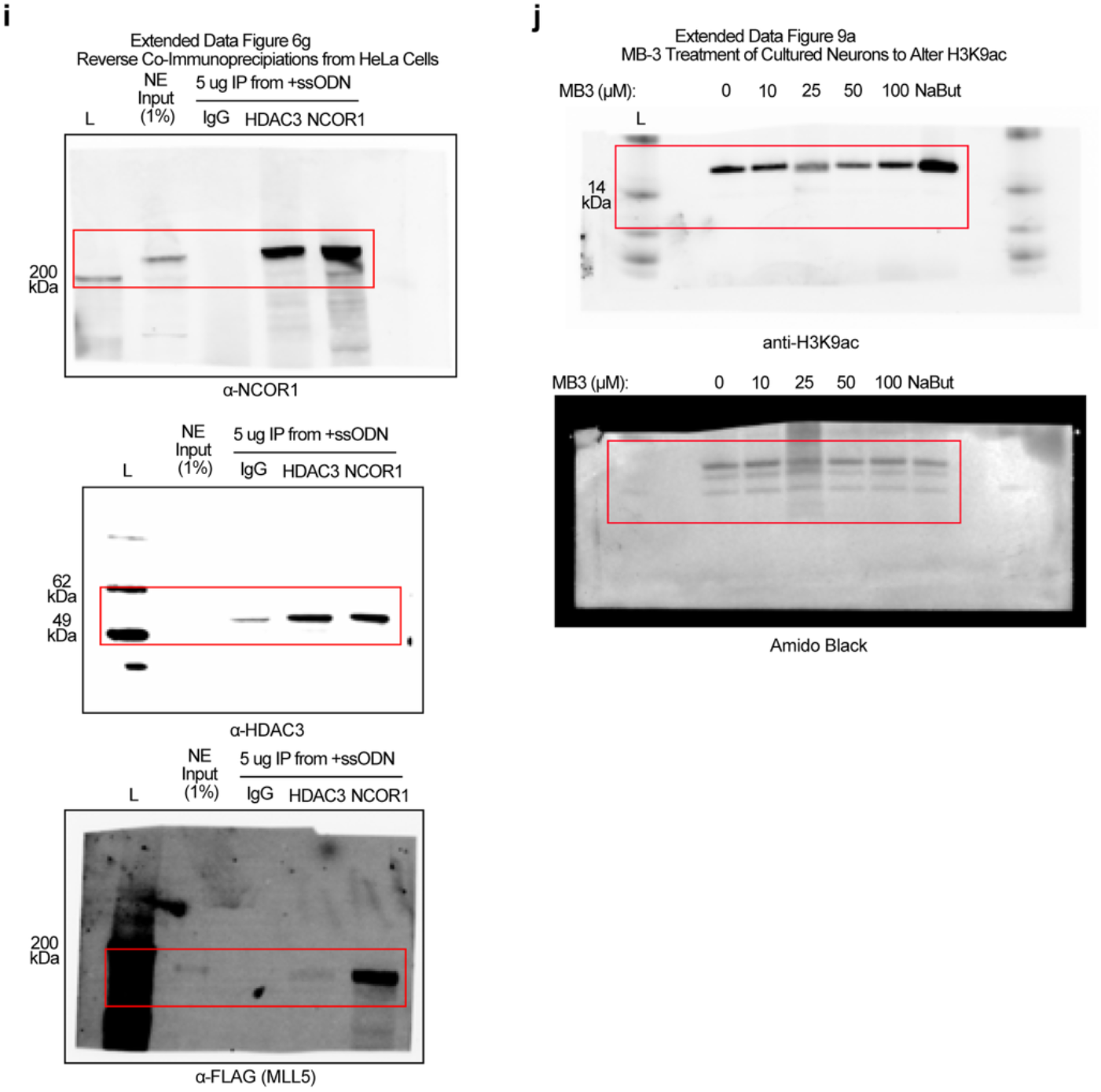
Uncropped Immunoblots. Uncropped immunoblots related to (**a**) Extended Data Fig. 2a, (**b**) Extended Data Fig. 2b, (**c**) Extended Data Fig. 6b, (**d**) Extended Data Fig. 6d, (**e**) Extended Data Fig. 6f, (**f-h**) Fig. 3b, (**i**) Extended Data Fig. 2g, (**j**) Extended Data Figure 9a. Red rectangles notate portion of blot included in figures, and blue rectangles notate portion of blots where analysis was performed but were not used as representative images.

## Supplementary Data

**Supplementary Table 1**: H3K4me3Q5ser broad vs. narrow peaks in E12.5 brain

**Supplementary Table 2**: DESeq2 table for WT vs. KO cerebellum RNA-seq

**Supplementary Table 3**: DESeq2 table for WT vs. KO mPFC RNA-seq

**Supplementary Table 4**: Differential splicing table for WT vs. KO rMATs analysis

**Supplementary Table 5**: LC-MS/MS analysis from FLAG-KMT2E immunoprecipitation

**Supplementary Table 6**: KMT2E-bound TSSs in cerebellum from CUT-RUN-sequencing

**Supplementary Table 7**: KMT2E-bound DEGs in cerebellum from CUT&RUN-sequencing

**Supplementary Table 8**: DESeq2 table for WT vs. HET from granule neuron RNA-seq

**Supplementary Table 9**: DESeq2 table for WT vs. KO from granule neuron RNA-seq

**Supplementary Table 10**: KMT2E peaks in granule neurons from CUT&RUN-sequencing

**Supplementary Table 11**: DESeq2 table for WT-Veh vs. HET-Veh from granule neuron RNA-seq

**Supplementary Table 12**: DESeq2 table for HET-MB-3 vs. HET-Veh from granule neuron RNA-seq

**Supplementary Table 13**: Clusters of WT-Veh vs. HET-Veh DEGs altered by MB-3

**Source Data File**

## Notes

### Competing Interest Statement

The authors have declared no competing interest.

